# Rho2 regulates granulocyte-triggered stress adaptation and cell wall remodeling in *Aspergillus fumigatus*

**DOI:** 10.1101/2025.03.10.642427

**Authors:** Dominik Ruf, Kristina Striegler, Sean Brazil, Hesham Elsaman, Kalpana Singh, Mathieu Lepas, Liyanage D. Fernando, Victor Brantl, Patricia Heß, Karl Dichtl, Vishukumar Aimanianda, Tuo Wang, Johannes Wagener

## Abstract

The airborne opportunistic fungal pathogen *Aspergillus fumigatus* poses a deadly threat to immunocompromised patients. Neutrophil granulocytes play a key role in the defense against invasive infections caused by this pathogen. The mechanisms by which *Aspergillus* defends itself against attacks by the immune system are only partially understood. Here we show that human granulocytes activate the cell wall integrity (CWI) pathway of *A. fumigatus* and that key components of the CWI such as the cell wall stress sensor MidA and the Rho GTPases Rho2 and Rho4 are important for the survival of *Aspergillus* hyphae under granulocyte attacks. A more detailed investigation of the role of Rho2 revealed that a mutant lacking *rho2* is less virulent in a *Galleria mellonella* infection model. Overexpression of Rho2 increases the resistance of *A. fumigatus* hyphae to killing by granulocytes. While a mutant lacking Rho2 has a normal cell wall composition, overexpression or constitutive activation of Rho2 leads to an altered cell wall composition and impairs growths of the pathogen. The fungicidal effect of constitutive activation of Rho2 signaling, which correlates with the formation of cell wall chitin bulges, depends on the CWI MAP kinase MpkA. However, Rho2 itself does not appear to be a direct activator of the CWI MAP kinase module. Our results support a model where Rho2 in *A. fumigatus* actively counteracts granulocyte attacks by upregulating cell wall biosynthesis, thereby strengthening the cell wall and aiding the fungus in surviving the stress condition.

## INTRODUCTION

*Aspergillus fumigatus* is an airborne opportunistic fungal pathogen. This mold is the most important species of the *Aspergillus* genus causing infection in humans. In immunocompromised individuals, *A. fumigatus* and other pathogenic aspergilli cause a severe invasive infection, called invasive aspergillosis, which is life-threatening and associated with a high case fatality rate (Kosmidis and Denning, 2015; Thompson and Young, 2021). Even in immunocompetent individuals, aspergilli may cause diseases, such as chronic pulmonary aspergillosis (CPA) or allergic bronchopulmonary aspergillosis (ABPA) (Kosmidis and Denning, 2015). The initial airborne infectious form of *Aspergillus*, the asexual spores (conidia), are primarily eliminated by alveolar macrophages or expelled by the mucociliary clearance. However, if this first line of defense fails, the conidia start to germinate forming hyphae, which subsequently penetrate the mucosa and invade the tissue (Thompson and Young, 2021; van de Veerdonk et al., 2017). Neutrophil granulocytes are the major players in the defense against invasive forms of *Aspergillus* spp. (van de Veerdonk et al., 2017). Congenial, the most important known risk factor for invasive aspergillosis is an abnormally low neutrophil granulocyte count in the blood, also known as neutropenia. In addition to neutropenia, many other conditions that affect granulocyte function significantly increase the risk for invasive aspergillosis, for example chronic granulomatous disease (CGD) or treatment with higher doses of corticosteroids (Kosmidis and Denning, 2015; van de Veerdonk et al., 2017).

It is well-established that neutrophil granulocytes are capable of killing *Aspergillus* hyphae both *in vivo* and *in vitro* (Gazendam et al., 2016; Ruf et al., 2018; van de Veerdonk et al., 2017). However, the mechanisms by which neutrophil granulocytes inactivate *Aspergillus* hyphae are only partially characterized. These mechanisms may involve the NADPH oxidase, reactive oxygen species (ROS), the release of proteins and peptides with antimicrobial activities from intracellular granules, extracellular antifungal vesicles, and potentially even neutrophil extracellular traps (NETs; under debate) (Gazendam et al., 2016; Shopova et al., 2020).

On the fungal side, several mechanisms have been identified that enable the pathogen to avoid being killed by immune cells. The underlying strategies can be divided roughly into two groups (reviewed in (van de Veerdonk et al., 2017)). The first group of mechanisms is based on preventing an antifungal response of the immune cells. This includes, in particular, certain properties of the fungal cell wall that contribute to the fungal organism not being recognized by the immune cells, thereby impeding the activation of antimicrobial defense mechanisms. Examples are the conidial rodlet layer, production of the polysaccharide galactosaminogalactan and synthesis of the cell wall carbohydrate ɑ-1,3-glucan, all of which were shown to mask pathogen-associated molecular patterns (PAMPs) (Hopke et al., 2018; Latgé and Chamilos, 2019; Wagener et al., 2020). The second group of mechanisms is the pathogen’s ability to adapt to the host environment. Although less understood, this includes mechanisms to withstand and adapt to the stress induced by the immune response (reviewed in (van de Veerdonk et al., 2017)). An important signaling pathway involved in the stress adaptation of fungi is the cell wall integrity (CWI) pathway (also known as cell wall salvage pathway). This pathway is best described in baker’s yeast and partially conserved in pathogenic fungi where it was shown to be important for virulence and tolerance to antifungal drugs (Dichtl et al., 2016). We and others have previously characterized parts of this pathway in *A. fumigatus* (Dichtl et al., 2016, 2012; Dirr et al., 2010; Jain et al., 2011; Rocha et al., 2016; Valiante et al., 2009). Two classes of conserved CWI pathway cell wall stress sensors are generally capable of detecting cell wall stress on the cell surface. These subsequently signal the stress via GTP exchange factors and Rho GTPases to a protein kinase C, which in turn activates a highly conserved mitogen-activated protein (MAP) kinase module that controls transcription factors (Fig. 1A) (Dichtl et al., 2016). While it is clear that certain parts of the CWI pathway contribute to antifungal resistance and to virulence, the involvement of this pathway in withstanding attempts by immune cells to kill the invading pathogen remains unclear (Dichtl et al., 2016; Valiante et al., 2015).

**Fig. 1.**
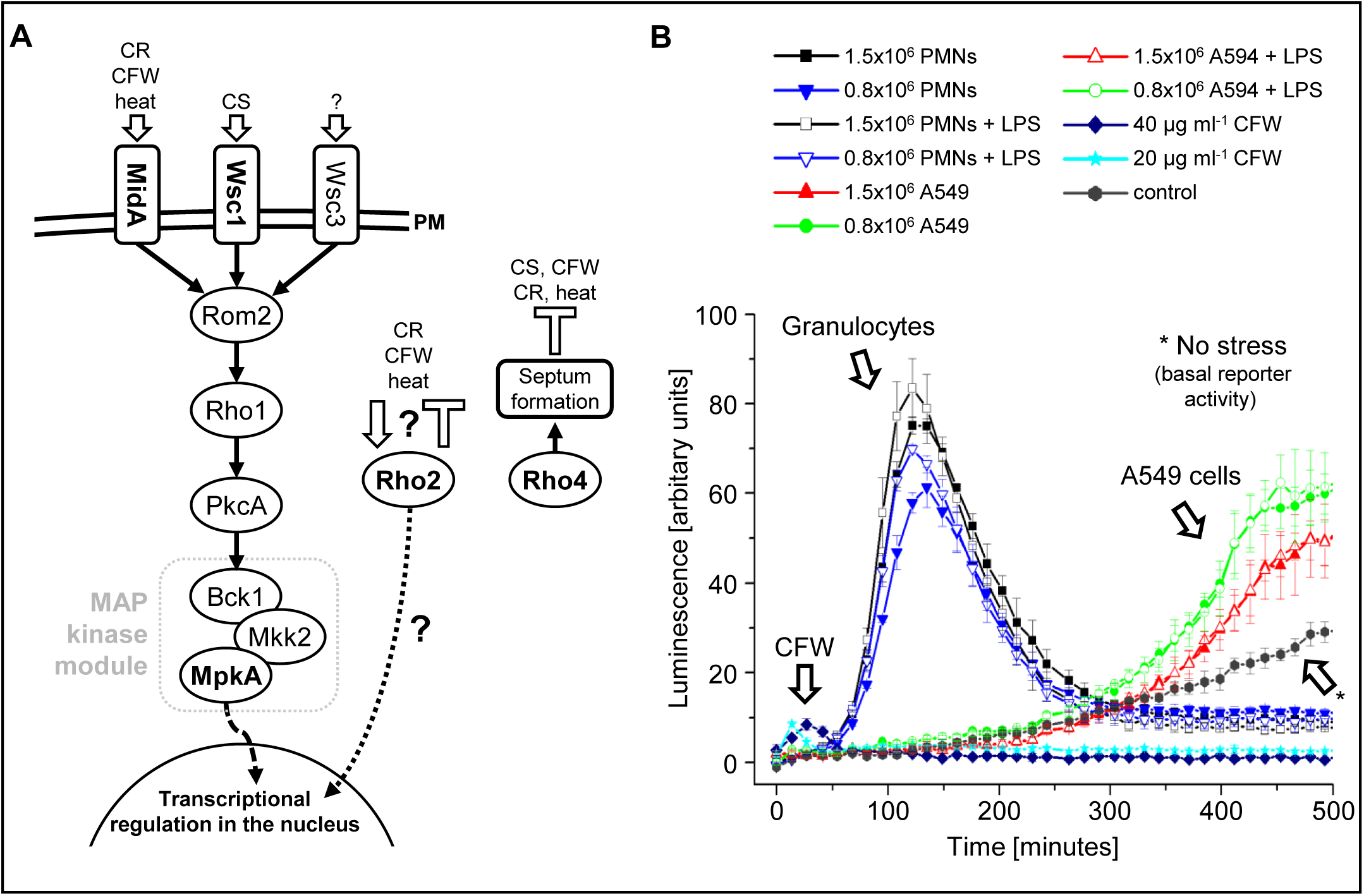
Human granulocytes trigger activation of the cell wall integrity pathway of *Aspergillus* hyphae. (A) Model of the cell wall integrity (CWI) stress signaling pathway of *A. fumigatus*. At least three plasma membrane (PM)-anchored cell wall stress sensors (MidA, Wsc1, and Wsc3) detect different types of cell wall stress, such as stress induced by Congo red (CR), calcofluor white (CFW), heat or caspofungin (CS). Upon detection of cell wall stress, these sensors activate Rho GTPase Rho1 via the GTP/GDP exchange factor Rom2. Rho1 then activates a protein kinase C (PkcA), which in turn activates a conserved mitogen-activated protein (MAP) kinase module (Bck1, Mkk2, MpkA). The MAP kinase MpkA regulates several transcription factors, including RlmA. The Rho GTPases Rho2 and Rho4 are also essential to withstand cell wall stress. While Rho4 is required for septum formation in *Aspergillus* hyphae, which is essential to protect the hyphal network against cell leakage, the exact functional role of Rho2 is unclear. (B) Conidia of an *A. fumigatus* cell wall stress reporter strain were inoculated in a 96-well plate in RPMI 1640 medium. After 10 h incubation at 37°C with 5% CO_2_, medium was supplemented with 0.5 mM luciferin and, when indicated, with indicated numbers of human granulocytes (PMNs) or A549 lung carcinoma epithelial cells, or the indicated concentration of CFW. When indicated, medium was additionally supplemented with lipopolysaccharides (200 ng/ml final concentration). Luciferase activity was measured over time while incubating at 37°C with 5% CO_2_. Data are representative of three independent experiments.

In this work, we aimed to clarify if the CWI pathway helps *A. fumigatus* hyphae to resist killing by immune cells. Using a novel killing assay (Ruf et al., 2018), we identified possible key players in the response of *A. fumigatus* hyphae to human granulocytes. We found that the Rho GTPases Rho2 and Rho4 and the CWI stress sensor MidA are important for survival of *A. fumigatus* hyphae that get attacked by granulocytes. We subsequently characterized the role of Rho2 and show that Rho2 regulates cell wall biogenesis specifically under stress conditions in a CWI MAP kinase-dependent and -independent way.

## RESULTS

### Human granulocytes trigger activation of the cell wall integrity pathway

We first aimed to clarify if neutrophils cause cell wall stress when they attack *Aspergillus* hyphae. We have previously established a luciferase-based *A. fumigatus* cell wall stress reporter strain (Geißel et al., 2018). This strain harbors a luciferase gene that is placed under the control of the *Aspergillus niger agsA* promoter, a well-characterized target of the conserved CWI transcription factor RlmA which is activated upon cell wall stress (Rocha et al., 2016). Hyphae of the reporter strain were exposed to human granulocytes or, as a control, to the chemical cell wall stressor calcofluor white, and the luciferase-dependent luminescence was monitored over time. As shown Fig. 1B, human granulocytes elicited a strong response from this reporter that began approximately 50-60 minutes after exposure and peaked at about 2 hours (Fig. 1B). Notably, a higher number of granulocytes yielded a stronger response of the reporter. Similarly, additional supplementation of medium with lipopolysaccharides (LPS) triggered a stronger response, which is in agreement with our previous observation that LPS stimulates the antifungal activity of granulocytes (Ruf et al., 2018). In comparison, a fungicidal concentration of the chemical cell wall stressor calcofluor white elicited an immediate response (Fig. 1B). This signal declined rapidly, presumably due to the imminent death of the exposed hyphae. In contrast to human granulocytes, a similar number of cells of the lung carcinoma cell line A549, a cell type known to phagocytose and inactivate *Aspergillus* conidia (Bertuzzi et al., 2014; 2020) but not hyphae, with or without LPS did not trigger a response within the first 350 minutes after exposure (Fig. 1B). A steady basal increase of the reporter signal was found in both the medium control and the control with the A549 cells, which probably reflects the exponential increase of the fungal biomass combined with a basal expression of the reporter over time. These data indicate that granulocytes cause cell wall stress in *A. fumigatus* hyphae that results in activation of the fungal CWI pathway.

### CWI stress sensor MidA, and Rho GTPases Rho2 and Rho4, but not Wsc1 contribute to survival of hyphae challenged by human granulocytes

We next asked whether previously identified determinants of the CWI of *A. fumigatus* (Dichtl et al., 2012) are important for *A. fumigatus* hyphae to withstand attacks from granulocytes. For this we used an assay that allowed us to quantify the killing activity of granulocytes against individual hyphae (Ruf et al., 2018). Four different CWI mutants were included in this approach. Two of these mutants lacked either the cell wall stress sensor MidA (Δ*midA*) or Wsc1 (Δ*wsc1*), both of which represent conserved activators of the CWI pathway that have different stress specificities (Dichtl et al., 2012). The other two mutants lacked either the Rho GTPase Rho2 (Δ*rho2*), which plays an important but currently undefined role in the CWI of *A. fumigatus* (Dichtl et al., 2012), or the Rho GTPase Rho4 (Δ*rho4*), which is required for septum formation in *Aspergillus* species (Dichtl et al., 2015, 2012). All these mutants showed wild-type-like germination and growth rates (Supplementary Fig. 1). As shown in Fig. 2, *A. fumigatus* hyphae that lack the cell wall stress sensor MidA or the Rho GTPases Rho2 or Rho4 were more efficiently killed by human granulocytes. In contrast, viability of the Δ*wsc1* mutant was similar to that of wild type *A. fumigatus* after exposure to human granulocytes. Reintroducing the deleted genes into the Δ*midA* and Δ*rho4* mutants restored the resistance to wild-type levels (Fig. 2). However, surprisingly, the complemented Δ*rho2* strain (*rho2*) was significantly more resistant to killing by human granulocytes than the wild type (Fig. 2).

**Fig. 2.**
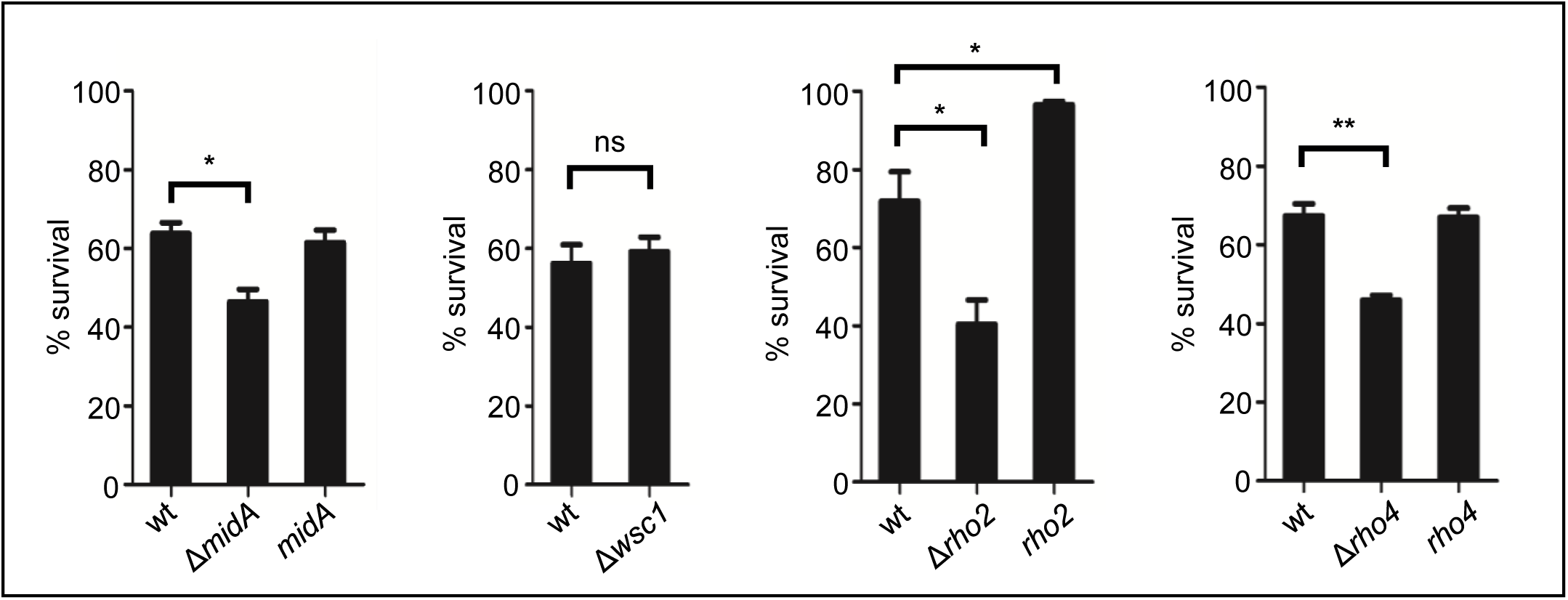
*A. fumigatus* hyphae lacking Rho2, MidA or Rho4 are more susceptible to killing by human granulocytes. Conidia of the indicated strains expressing mitochondria-targeted GFP were inoculated in RPMI-1640. After 9.5 h incubation at 37°C with 5% CO_2_, granulocytes were added to the formed hyphae . After 2 h co-incubation at 37°C with 5% CO_2_, the ratio of vital hyphae was analyzed as described in the material and method section. The data depicted in the graphs are based on the results of at least 3 independent experiments for each graph. Statistical significance (**, p ≤ 0.01; *, p ≤ 0.05; ns, not significant) was calculated with the one-way analysis of variance (ANOVA) with Tukey’s multiple comparison post-test when comparing more than two groups or a two-tailed unpaired (assuming unequal variances) Student’s *t*-test when comparing two groups. The error bars indicate standard deviations.

### Overexpression of Rho2 protects from killing by neutrophil granulocytes

The Rho GTPase Rho4 has an established role in septum formation; we and others have previously shown that septum formation significantly contributes to survival under cell wall stress conditions by compartmentalizing hyphae (Dichtl et al., 2015; Geißel et al., 2018; Souza et al., 2021). MidA is a cell wall stress sensor and known activator of the canonical CWI pathway, which is implicated in the virulence of multiple fungal pathogens (Dichtl et al., 2016, 2012). Little was known about the function of Rho2, and we therefore aimed to investigate its role further. The overcompensation of defects in a complemented Δ*rho2* mutant could be explained by an increased expression of the re-introduced gene. The Δ*rho2* mutant was complemented by transformation of the strain with a circular plasmid containing the promoter region and the coding sequence of *rho2*. Due to the strategy used in this work to construct mutants (use of a non-homologous end joining-deficient strain, circular plasmids for complementation), it is possible that the plasmid has integrated multiple times at defined loci in the genome that share homologous sequences with the plasmid. Further, the lack of the 3’ untranslated region of the *rho2* gene in the complementation plasmid may have influenced the expression. An analysis of the integration sites and frequency of integration using a method based on integration site-specific polymerase chain reactions (PCRs) revealed that the complementation plasmid had integrated more than once in the genome of the Δ*rho2* mutant. In agreement with these findings and the suspected overcompensation of the defects in the Δ*rho2* mutant, growth tests on solid media showed that the complemented Δ*rho2* strain is more resistant than the wild type to the cell wall stressor calcofluor white (Supplementary Fig. 2A). We therefore aimed to construct an additional complemented Δ*rho2* strain which harbors only a single copy of the *rho2* gene. A new plasmid was constructed that additionally harbored parts of the 3’ untranslated region of *rho2*. Transformation of the Δ*rho2* mutant resulted in clones (*rho2_add3’_*) with one (#8-11) or more than one integration (#1-3,5-7) of the plasmid (Supplementary Fig. 2B and C). As shown in Supplementary Fig. 2A, multiple integrations of the new plasmid also resulted in increased calcofluor white resistance, while integration of only a single copy resulted in a tolerance similar to that of wild type.

To confirm the *rho2* expression-dependent stress tolerance of *A. fumigatus*, we constructed conditional *rho2* mutants (*rho2_tetOn_*) by replacing the endogenous promoter of *rho2* with a doxycycline-dependent Tet-On promoter. Under repressed conditions, these mutants exhibited an increased susceptibility to calcofluor white similar to that of the Δ*rho2* mutant (Fig. 3A). Under induced conditions, the *rho2_tetOn_* mutants were more resistant to calcofluor white (Fig. 3A). Analysis of the susceptibility of a *rho2_tetOn_* mutant to killing by human granulocytes demonstrated that the *rho2_tetOn_* mutant was more resistant under induced conditions, while it showed an increased susceptibility under repressed conditions (Fig. 3B).

**Fig. 3.**
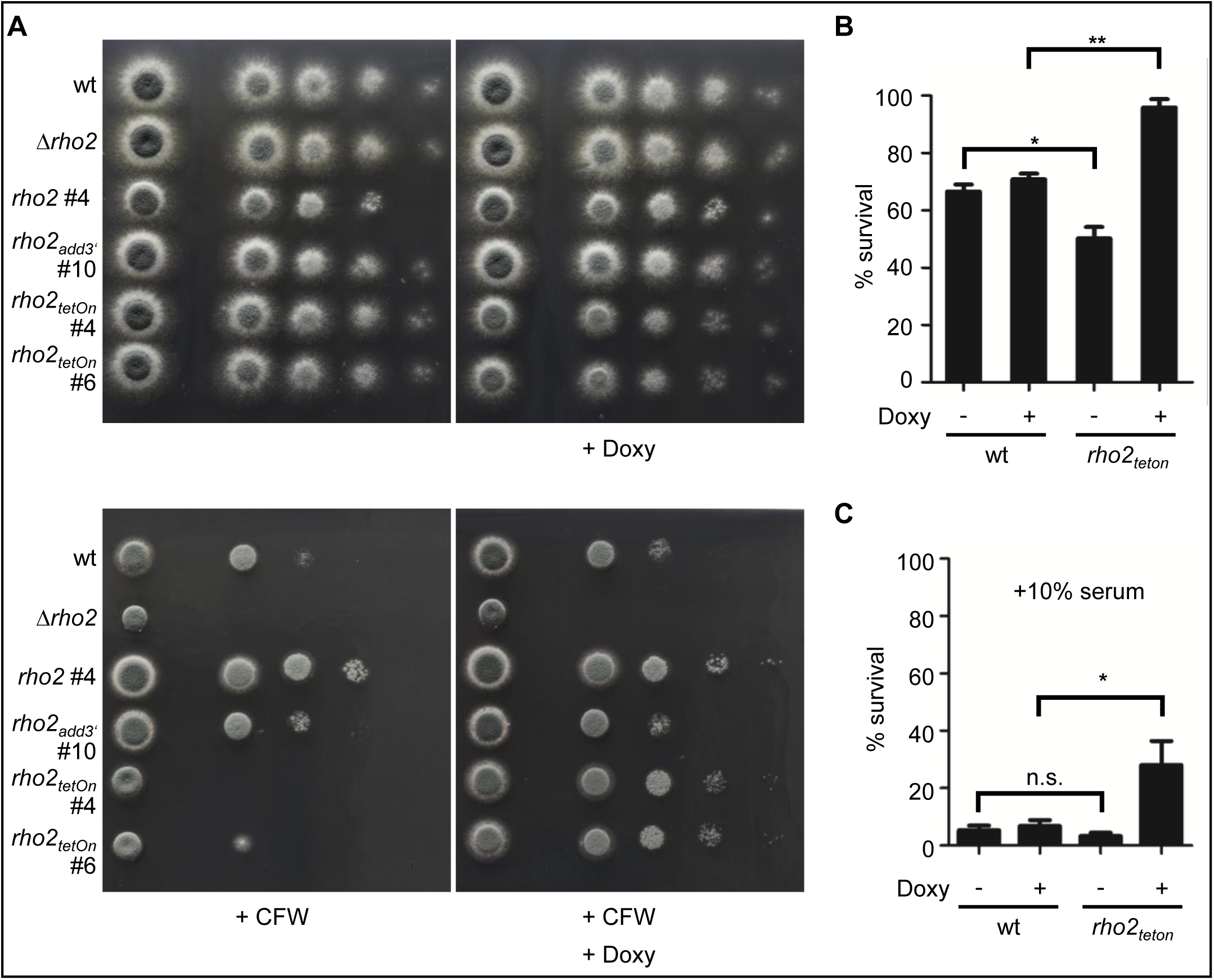
Overexpression of Rho2 increases the resistance to cell wall perturbing conditions and killing by human granulocytes. (A) In a series of 10-fold dilutions derived from a starting suspension of 5 × 10^7^ conidia ml^-1^ of the indicated strains, aliquots of 3 µl were spotted on AMM agar plates. The numbers (#) indicates the respective clone number. When indicated, medium was supplemented with 7.5 µg ml^-1^ doxycycline (+Doxy) or 25 µg ml^-1^ calcofluor white (CFW). Representative images were taken after 34h incubation at 37°C. (B and C) Conidia of the indicated strains expressing mitochondria-targeted GFP were inoculated in RPMI-1640. When indicated (+Doxy), medium was supplemented with 7.5 µg ml^-1^ doxycycline. After 9.5 h incubation at 37°C with 5% CO_2_, granulocytes were added. After 2 h co-incubation at 37°C with 5% CO_2_, the ratio of vital hyphae was analyzed. The data depicted in the graphs are based on the results of at least 3 independent experiments for each graph. Statistical significance (**, p ≤ 0.01; *, p ≤ 0.05; n.s., not significant) was calculated with the one-way analysis of variance (ANOVA) with Tukey’s multiple comparison post-test. The error bars indicate standard deviations.

Both the Δ*rho2* strain complemented with multiple copies of *rho2* and the conditional *rho2_tetOn_* mutant under induced conditions appear to be insensitive to killing by human granulocytes under the chosen experimental conditions. We previously found that the presence of serum increases the antifungal activity of human granulocytes (Ruf et al., 2018). Therefore, we analyzed the susceptibility of *A. fumigatus* to killing by human granulocytes in the presence of 10% (v/v) serum in the medium. As shown in Fig. 3C, the increased resistance was also evident for the induced conditional *rho2_tetOn_* mutant; nonetheless, the presence of serum enabled the granulocytes to inactivate a significant number of the *rho2_tetOn_* mutant hyphae under induced conditions (Fig. 3C). Taken together, these results demonstrate that Rho2 is important for *A. fumigatus* hyphae to withstand killing by granulocytes and that an increased expression of rho2 can even increase the resistance beyond that of wild type.

### Rho2 localizes to the plasma membrane

We had previously reported that Rho2 shows cytosolic distribution without a pronounced accumulation pattern at any subcellular structures (Dichtl et al., 2012). When constructing the conditional *rho2_tetOn_* mutant, we found that the annotation of the *rho2* locus in the reference genome database had changed. The start codon is now predicted at a position 192 base pairs upstream of the previously annotated position, while the previously annotated start codon is now located within the first intron’s sequence. The inaccurate annotation of the start codon led to an incorrect GFP-Rho2 hybrid protein expression construct used for determining the subcellular localisation in our previous work (Dichtl et al., 2012). To determine the correct subcellular localisation of Rho2, a corrected GFP-Rho2 hybrid protein expression construct was cloned and transformed into *A. fumigatus*. As shown in Fig. 4A and B, the GFP-Rho2 hybrid protein was now primarily localized at the plasma membrane and septa of *A. fumigatus* hyphae.

**Fig. 4.**
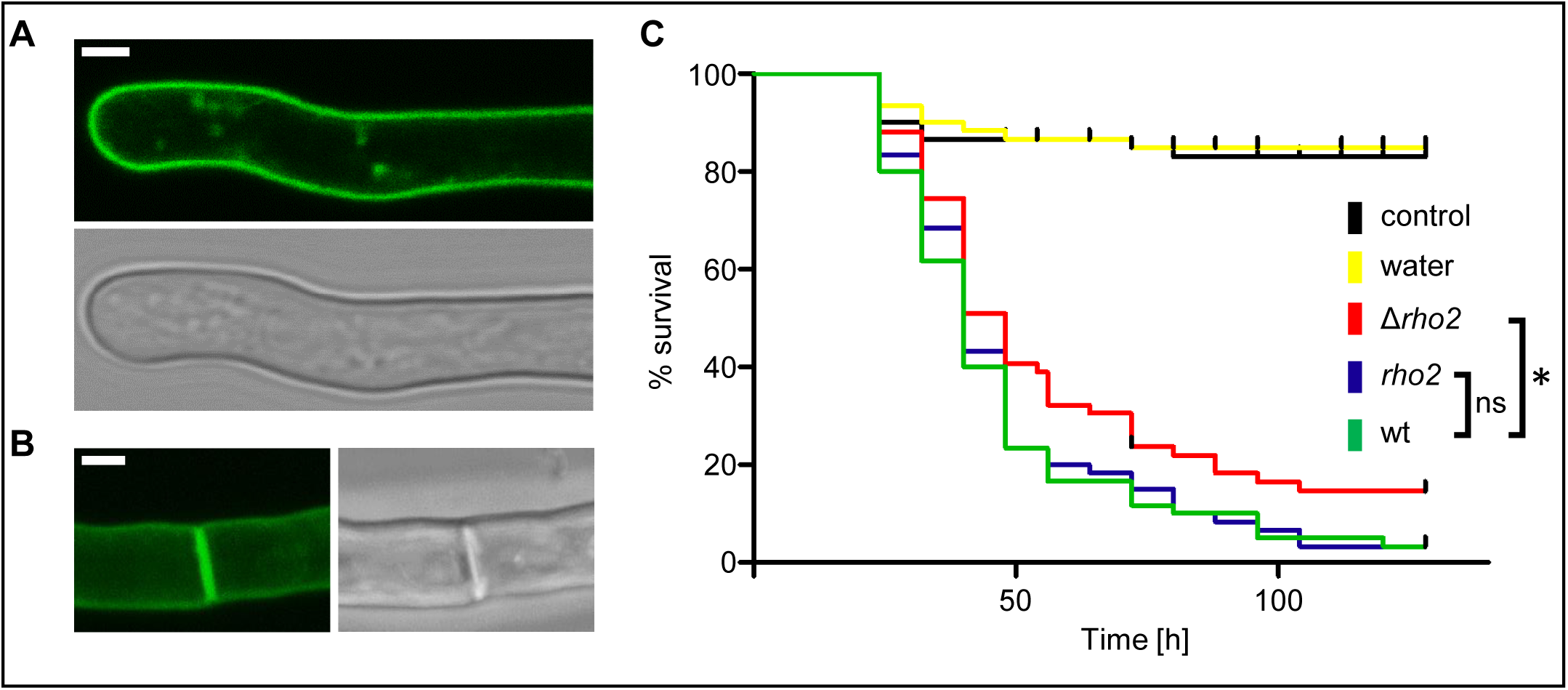
Rho2 localizes to the cell membrane and septa and contributes to virulence in a *G. mellonella* infection model. (A and B) Conidia of a wild-type strain expressing an N-terminally GFP-tagged Rho2 were inoculated in AMM medium and incubated for 14 h at 37°C. Fluorescence (A, top image; B, left image) and bright-field (A, bottom image; B, right image) images were taken with a confocal microscope and represent one optical section (A) of a living hypha or (B) a z-stack of optical sections covering the entire living hypha in focus. The bar represents 2 µm. (C) Larvae of *G. mellonella* were injected with conidia suspensions of the indicated stains (5 × 10^5^ conidia in 10 µl water per larva). Larvae that were either not injected (untreated) or injected with 10 µl ddH_2_O served as control groups. Larvae were subsequently kept in the dark at 37°C and examined every 8 h. The cumulative survival in percent of the larvae of 3 experiments (20 larvae per condition) was plotted in the Kaplan-Meier plot. Significance was calculated with a log-rank (Mantel-Cox) test (*, p ≤ 0.05; ns, not significant).

### Lack of Rho2 results in reduced virulence in a *Galleria mellonella* infection model

The Δ*rho2* deletion mutant is more sensitive to cell wall stress and showed an increased susceptibility to killing by granulocytes. On the other hand, the Δ*rho2* strain complemented with multiple copies of *rho2* was more resistant. We therefore asked whether virulence of these two strains is altered in an animal infection model. The virulence of the Δ*rho2* mutant and of the Δ*rho2* strain complemented with multiple copies of *rho2* was analyzed and compared to wild type in a *Galleria mellonella* infection model. Even though the Δ*rho2* strain complemented with multiple copies of *rho2* was more resistant to killing by granulocytes and the stressor calcofluor white, its virulence was not significantly different from that of wild type (Fig. 4C). However, deletion of *rho2* resulted in a moderate but significant decrease in virulence compared to wild type (Fig. 4C).

### Constitutive activation of Rho2 is lethal

Next, we asked what regulatory role of Rho2 might have facilitated the increased resistance to calcofluor white-induced and granulocyte-triggered cell wall stress. Overexpression of *rho2* resulted in significantly decreased radial growth which was most prominent on complex media (Supplementary Fig. 3). In order to investigate the effects presumably triggered by overexpression of Rho2, we decided to construct a conditional *rho2* mutant which has inserted a point mutation in a conserved region of the enzyme which had been shown to result in constitutive activation of similar Rho-GTPases in other fungal species (*rho2*^G20V^*_tetOn_*) (Calonge et al., 2003; Hirata et al., 1998). As shown in Fig. 5A and B, conditional expression of a constitutively active *rho2* allele completely suppressed the growth of *A. fumigatus*. At this point it remained unclear whether the constitutive activation of Rho2 leads to a complete arrest of growth only or it even has a fungicidal effect over time. To clarify if constitutive activation of Rho2 is lethal in *A. fumigatus*, we analyzed the impact of *rho2*^G20V^ expression on the viability using the mitoFLARE assay (Ruf et al., 2018). Conidia of the *rho2*^G20V^*_tetOn_* mutant expressing a mitochondria-targeted green fluorescent protein (mito-GFP) were cultured under repressed conditions to form short hyphae. Subsequently, *rho2*^G20V^ expression was induced in the hyphae, and the mitochondrial morphology and dynamics were analyzed over time with a spinning disc confocal microscope (Fig. 5C). Expression of *rho2*^G20V^ resulted in a rapid arrest of hyphal growth (Supplementary video 1 and 2). Over time, the viability of the hyphae decreased (Fig. 5D). Interestingly, it took almost two days for more than 50% of the hyphae to die. After three days, approximately 75% of the hyphae were dead (Fig. 5D).

**Fig. 5.**
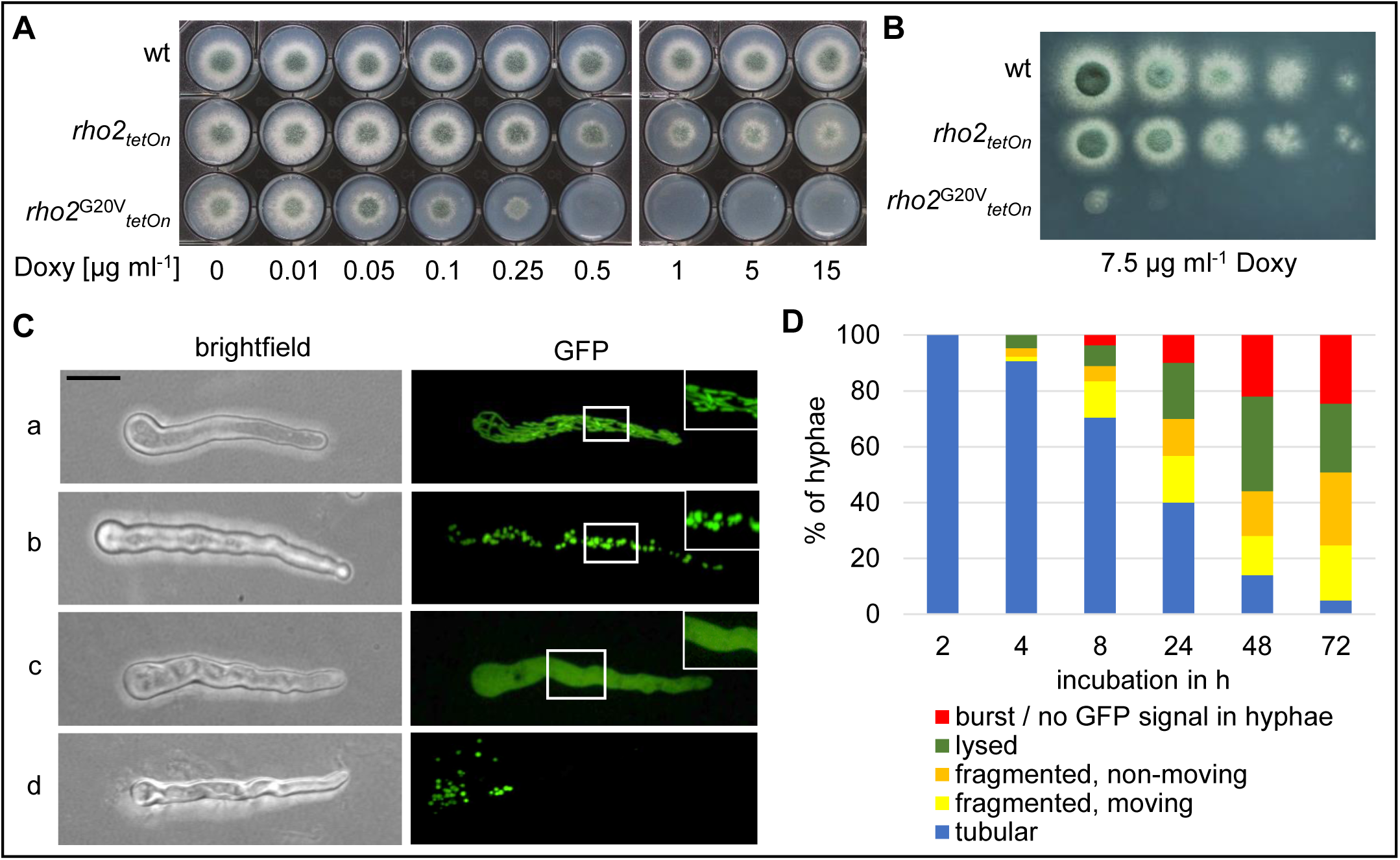
Expression of constitutively active Rho2 is lethal. (A) From suspensions of 5 × 10^7^ conidia ml^-1^ of the indicated strains, aliquots of 3 µl (1,500 conidia) were inoculated on AMM agar supplemented with the indicated amount of doxycycline (Doxy) and incubated at 37°C for 35 h. (B) In a series of 10-fold dilutions derived from a starting suspension of 5 × 10^7^ conidia ml^-1^ of the indicated strains, aliquots of 3 µl were spotted on AMM agar plates. When indicated (+Doxy), medium was supplemented with 7.5 µg ml^-1^ doxycycline. After 35-42 h incubation at 37°C, representative images were taken. (C and D) The viability of hyphae expressing *rho*^G20V^ was analyzed with the mitoFLARE assay. Conidia of the *rho2*^G20V^*_tetOn_* mutant expressing mitochondria-targeted GFP were inoculated in AMM. After 8 h incubation at 37°C, medium was supplemented with 7.5 µg ml^-1^ doxycycline, followed by incubation at 37°C. The mitochondrial morphology and dynamics of hyphae were analyzed with a spinning disc confocal fluorescence microscope at different time points. (C) Representative bright-field (left panels) and fluorescence (right panels) images of hyphae with tubular (a) or fragmented (b) mitochondrial morphology, lysed mitochondria (c) or of a hypha after burst are shown. Fluorescence images represent optical stacks covering the entire hyphae in focus. The bar represents 10 µm and is applicable to all subpanels. (D) The proportion of hyphae with the indicated mitochondrial morphology and dynamics patterns after the indicated incubation time in the presence of doxycycline is shown in the graph. At least 50 hyphae were analyzed at each time point.

### Expression of constitutively active Rho2^G20V^ causes abnormal cell wall synthesis

We speculated that the expression of constitutively active Rho2^G20V^ may affect the cell wall of *A. fumigatus*. A major component of the fungal cell wall is chitin which can be readily stained with the chitin-specific dye calcofluor white. As shown in Fig. 6A, expression of the constitutively active Rho2^G20V^ resulted in patchy calcofluor white-stainable cell wall bulges extending into the internal volume of the *Aspergillus* hyphae. The formation of these bulges correlated with an overall significant increase of cell wall chitin when compared to wild type (Fig. 6B). The patchy cell wall bulges combined with the subsequent death of the Rho2^G20V^- expressing hyphae were reminiscent of the cell wall patches formed upon treatment of *A. fumigatus* with azole antifungals through inhibition of the 14α-demethylase (CYP51) (Geißel et al., 2018). However, the morphologic appearance of the Rho2^G20V^-triggered bulges appeared much more rounded and less rough compared to the azole-induced cell wall carbohydrate patches (Fig. 6C). We wondered whether Rho2 could be an important factor for the azole-induced cell wall carbohydrate patches. As shown in Fig. 6D, the Δ*rho2* deletion mutant was also capable of forming cell wall carbohydrate patches upon treatment with azole antifungals. These results suggest that Rho2 controls cell wall biogenesis, but that it is not required for the azole-induced formation of cell wall carbohydrate patches in *A. fumigatus*.

**Fig. 6.**
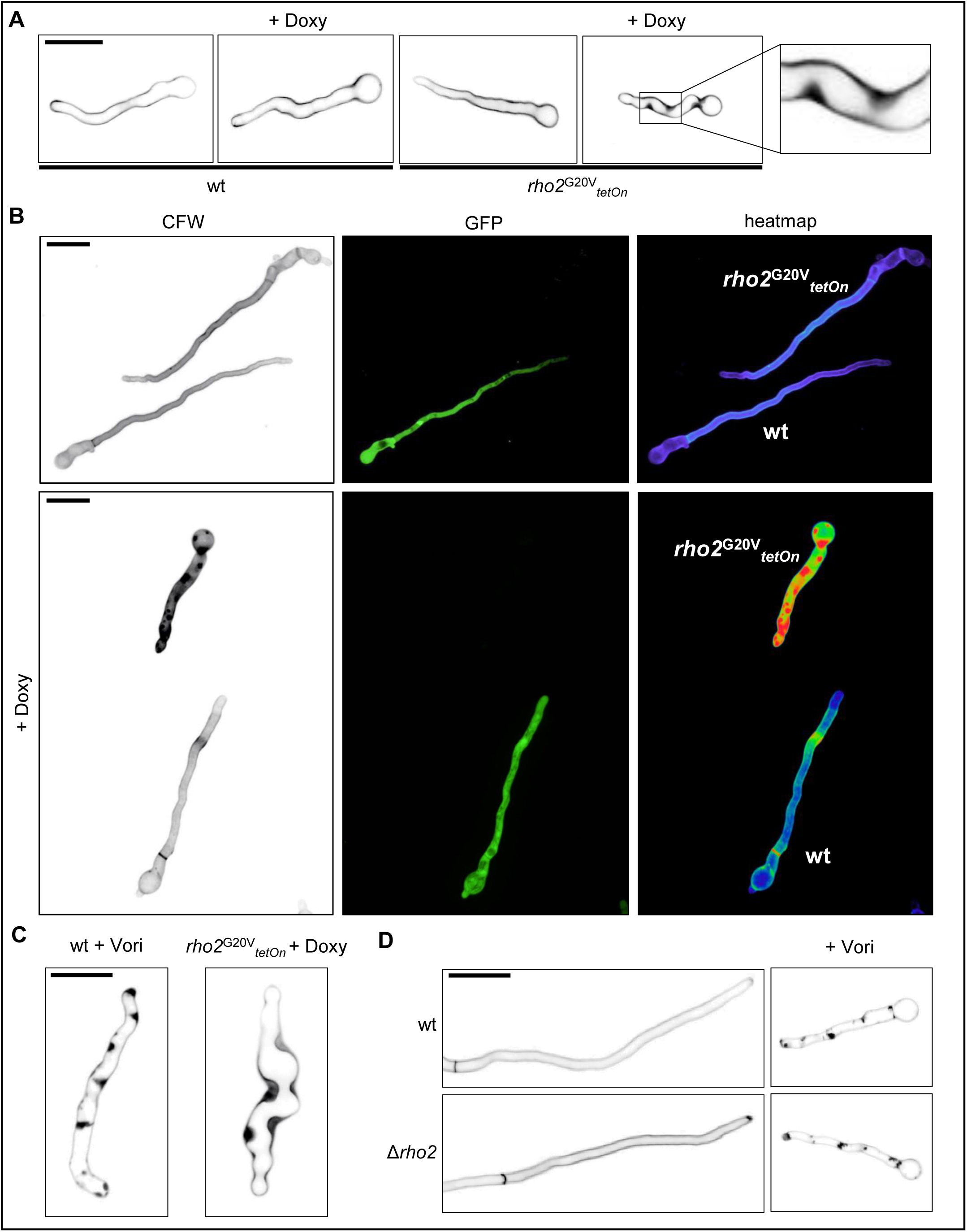
Expression of constitutively active Rho2 triggers formation of patchy cell wall bulges. (A and B) Conidia of *rho2*^G20V^*_tetOn_* were inoculated in AMM and incubated at 37°C. The experiment depicted in (B) was additionally inoculated with conidia of wild type expressing cytosolic GFP (wt). When indicated (+Doxy) medium was supplemented with 7.5 µg ml^-1^ doxycycline after 8 h incubation. (C and D) The indicated strains were inoculated in AMM and incubated at 37°C. When indicated, medium was supplemented after approximately 9 h incubation with 7.5 µg ml^-1^ doxycycline (+Doxy) or 1.27 µg ml^-1^ voriconazole (+Vori). (A - D) After a total incubation of 10h (A), 12h (B), or 15h (C and D) samples were stained with calcofluor white and analyzed with a spinning disc confocal fluorescence microscope. Representative images of the calcofluor white (chitin) fluorescence are depicted (A, left panels in B, C and D). In addition, representative images of the GFP fluorescence and images of the fluorescence intensity of calcofluor white in false color are shown in the middle and right panels in (B), respectively. (A - D) The depicted images represent optical stacks of the entire hyphae in focus. Bars represent 10 μm and are applicable to all subpanels.

### Lethality of constitutively active Rho2 and Rho2-triggered cell wall remodeling partially depends on the CWI MAPK kinase

The CWI pathway in fungi is thought to primarily involve the conserved Rho GTPase Rho1, which activates the conserved protein kinase C that in turn activates a conserved MAP kinase module (Fig. 1A) (Dichtl et al., 2016). It remained unclear which role *A. fumigatus* Rho2 and its homologs in other fungal species could have in this signaling cascade. If Rho2 is an activator of the canonical CWI pathway, expression of Rho2^G20V^ should trigger activation of the luciferase-based cell wall stress reporter construct. As shown in Fig. 7A, expression of *rho2*^G20V^ resulted in a strong and steadily increasing response from this reporter over time. However, while a fungicidal concentration of the chemical cell wall stressor calcofluor white triggered a luciferase signal in less than 20 minutes, followed by a rapid decline after 80 minutes presumably due to fungal death, the response triggered by expression of *rho2*^G20V^ became noticeable approximately one hour after adding doxycycline (Fig. 7A). In agreement with our previous findings, exposure to azoles, which trigger the formation of fungicidal cell wall carbohydrate patches (Geißel et al., 2018), caused a dramatic response of the report after approx. 3-4 hours. Interestingly, this azole-induced response quickly outpaced the response to expression of *rho2*^G20V^ (Fig. 7A). This suggests that strong activation of Rho2 over time can lead to activation of the CWI pathway.

**Fig. 7.**
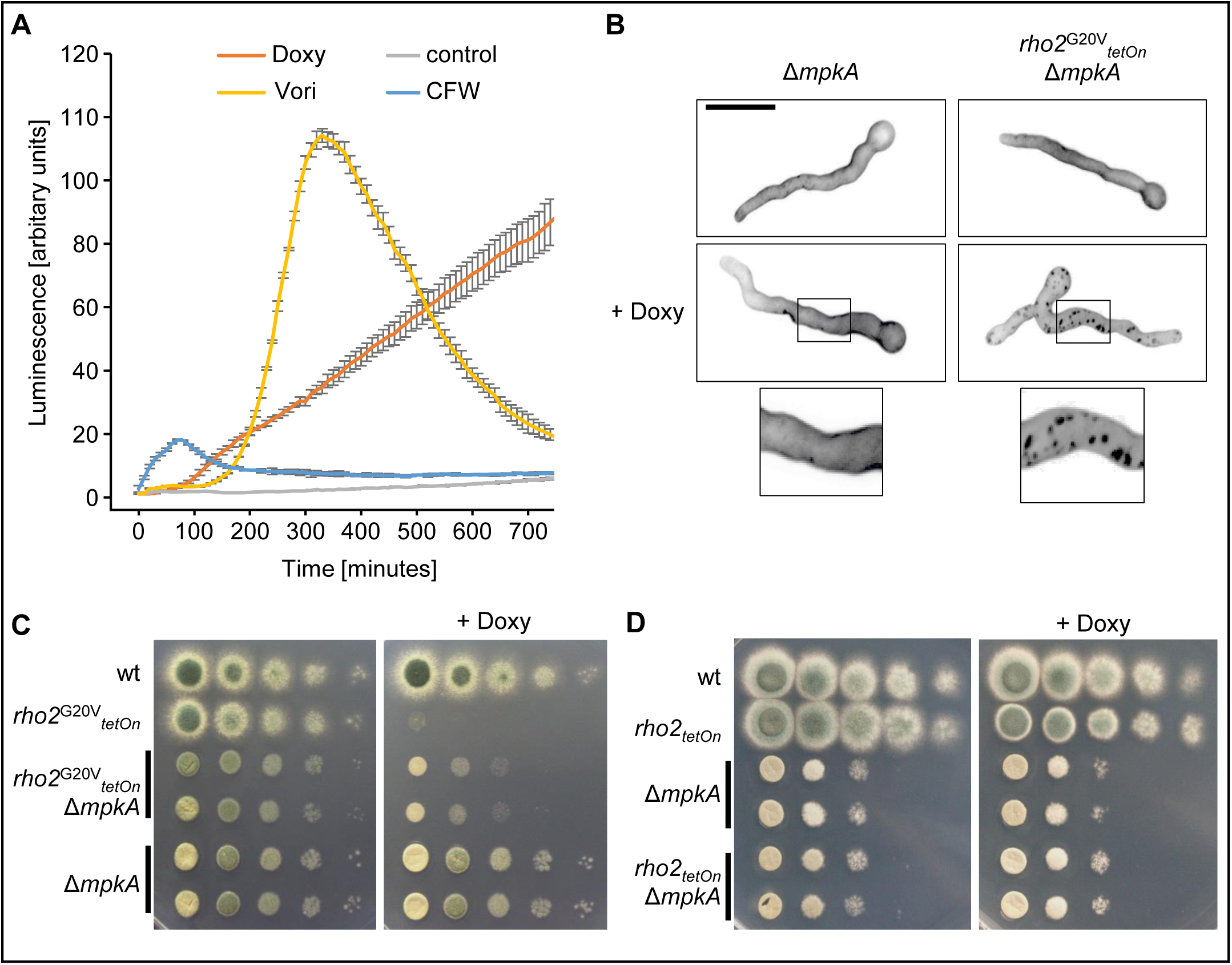
MAP kinase MpkA contributes to the toxicity of constitutively active Rho2. Conidia of the luciferase-based *rho2*^G20V^*_tetOn_* cell wall stress reporter strain were inoculated in a 96-well plate in RPMI 1640/MOPS medium. After 7 h incubation at 37°C, medium was supplemented with 0.5 mM luciferin and, when indicated, with 50 µg ml^-1^ calcofluor white (CFW), 2 µg ml^-1^ voriconazole (Vori), 25 µg ml^-1^ doxycycline (Doxy), or with no drug (control). Luciferase activity was measured over time while incubating at 37°C. Data are representative of three independent experiments. (B) Conidia of the indicated strains were inoculated in AMM and incubated at 30°C. When indicated (+Doxy) medium was supplemented with 7.5 µg ml^-1^ doxycycline after 12 h incubation. After a total incubation of 16 h samples were stained with CFW and analyzed with a spinning disc confocal fluorescence microscope. Depicted are representative images of optical stacks of the calcofluor white fluorescence (chitin) that cover the entire hyphae in focus. The bar represents 10 μm and is applicable to all subpanels. (C and D) In a series of 10-fold dilutions derived from a starting suspension of 5 × 10^7^ conidia ml^-1^ of the indicated strains, aliquots of 3 µl were spotted on AMM agar plates. Two independent clones of *rho2*^G20V^*_tetOn_* Δ*mpkA* (C), Δ*mpkA* (C and D) and *rho2_tetOn_* Δ*mpkA* (D) were included in the analysis. When indicated, medium was supplemented with 7.5 µg ml^-1^ doxycycline (+Doxy). Representative images were taken after 45 h incubation at 30°C or 42 h incubation at 40°C.

We also asked whether the cell wall bulges that get formed upon expression of the constitutively active Rho2^G20V^ depend on activation of the canonical CWI pathway. For this purpose, mutants were constructed in which the gene coding for MpkA, the central MAP kinase of the canonical CWI MAP kinase module (Fig. 1A), was deleted (Δ*mpkA*). Interestingly, as shown in Fig. 7B, the conditional *rho2*^G20V^*_tetOn_* Δ*mpkA* mutant was able to form cell wall patches when the constitutively active Rho2^G20V^ was expressed. But these patches appeared to be generally smaller, more numerous and spottier when compared to those observed with the *rho2*^G20V^*_tetOn_* single mutant in previous experiments. Consequently, while the formation of the Rho2^G20V^-triggered cell wall patches does not essentially depend on MpkA, the MAP kinase module of the CWI pathway appears to enhance the markedness of the bulges.

We then assessed the importance of MpkA for the toxicity of Rho2^G20V^ expression. As shown in Fig. 7C, deleting *mpkA* enabled the conditional *rho2*^G20V^*_tetOn_* mutant to grow and survive under induced conditions. At the same time, the growth rates were still reduced compared to the Δ*mpkA* deletion mutant (Fig. 7C). Interestingly, overexpression of the non-mutated wild-type Rho2 did not improve the growth of Δ*mpkA* deletion mutant under heat stress (Fig. 7D). Taken together, these data demonstrate that the toxicity of constitutively active Rho2 requires a functional CWI MAP kinase module. However, Rho2 does not appear to be a strong direct activator of the MAP kinase module.

### Impact of Rho2 on cell wall composition

Our results suggested that Rho2 modulates cell wall biogenesis. To better understand the putative role of Rho2 in cell wall biogenesis, we analyzed the cell wall of wild type and the conditional *rho2_tetOn_* mutant under repressed and induced conditions. The intact fungal materials were directly used for measuring the one-dimensional (1D) ^13^C solid-state Nuclear Magnetic Resonance (ssNMR) using a ^1^H-^13^C cross polarization experiment that selectively detects the rigid carbohydrates and protein/lipid components (Fig. 8A). Interestingly, the conditional *rho2_tetOn_* mutant under repressed conditions, despite being highly susceptible to cell wall stress and killing by granulocytes, showed a cell wall composition very similar to the wild type (Fig. 8A and B). In contrast, overexpression of Rho2 by culturing the *rho2_tetOn_* mutant with doxycycline resulted in a marked change in peak intensities representative of different cell wall polysaccharides when compared to the wild type with doxycycline (Fig. 8A and B, Supplementary Fig. 4). A minor change in the lipid composition is also noted as revealed by the decline of intensities at 31 and 33 ppm, two peaks belonging to the acyl chains of lipids (Fig. 8A). The rigid polysaccharide core of the *A. fumigatus* cell wall was recently identified to predominantly contain chitin, α-1,3-glucan, and β-1,3-glucan, with the latter two also distributed within the flexible domains of the cell wall (Chakraborty et al., 2021; Kang et al., 2018a). Relative to β-1,3-glucan, the chitin content in the rigid fraction of *rho2_tetOn_* mutant under induced conditions showed a slight increase of less than 10%, whereas the amount of α-1,3-glucan notably by approximately 50%. In the inner core of the *A. fumigatus* cell wall, β-glucan is the best-hydrated molecule, contrasting with the rigid and hydrophobic nature of chitin and α-1,3-glucan, which are physically packed together (Kang et al., 2018b). Therefore, the cumulative effect of these observed changes enhanced the rigidity of the cell wall. The slight increase in chitin content was observed through the higher intensity in the chitin carbon 4 (Ch4) peak and a slight increase in intensity for the mixed carbon 1 peaks of chitin and β-glucan (Ch1/B1 peak), with individual β-glucan peaks like B3 and B5 remaining unchanged. The increase in α-1,3-glucan was detected through the significantly higher intensities observed in the A1, A3, and A2/5 peaks. These observations were highly reproducible across three replicates (Supplementary Fig. 4). We also analyzed the cell walls of wild type and the conditional *rho2_tetOn_* mutant under repressed and induced conditions chemically (Feng et al., 2011). As shown in Fig. 8C, the ratio of the alkali-soluble to alkali-insoluble fraction of the cell wall in the induced *rho2_tetOn_* mutant was shifted significantly towards the alkali-soluble fraction. Furthermore, a relative decrease of glucose in the alkali-insoluble fraction (representing β-1,3-glucan) and a relative increase of glucose in the alkali-soluble fraction (primarily representing α-1,3-glucan) were noticed upon overexpression of Rho2 (Fig. 8D), both confirming ssNMR observations. Surprisingly, a slight decrease of glucosamine was observed in the alkali-insoluble fraction upon overexpression of Rho2, suggesting a relative decrease of cell wall chitin. At first glance, this finding appears to contradict the ssNMR results. However, the different results could be linked to the differences in the methods used. The chemical analysis only detects chitin tightly bound to the cell wall (Gow and Lenardon, 2023), while the specific ssNMR technique employed here primarily detects the physically rigid and structurally important molecules within the cell wall (Latgé and Wang, 2022). Hence, the results suggested a relative decrease of cell wall-linked chitin with a concomitant relative increase of free chitin. Taken together, our NMR and chemical results demonstrate that Rho2 can trigger a reshape of the cell wall which affects the relative abundance and rigidity of β-1,3-glucan, α-1,3-glucan and chitin, most likely resulting in a more rigid cell wall.

**Fig. 8.**
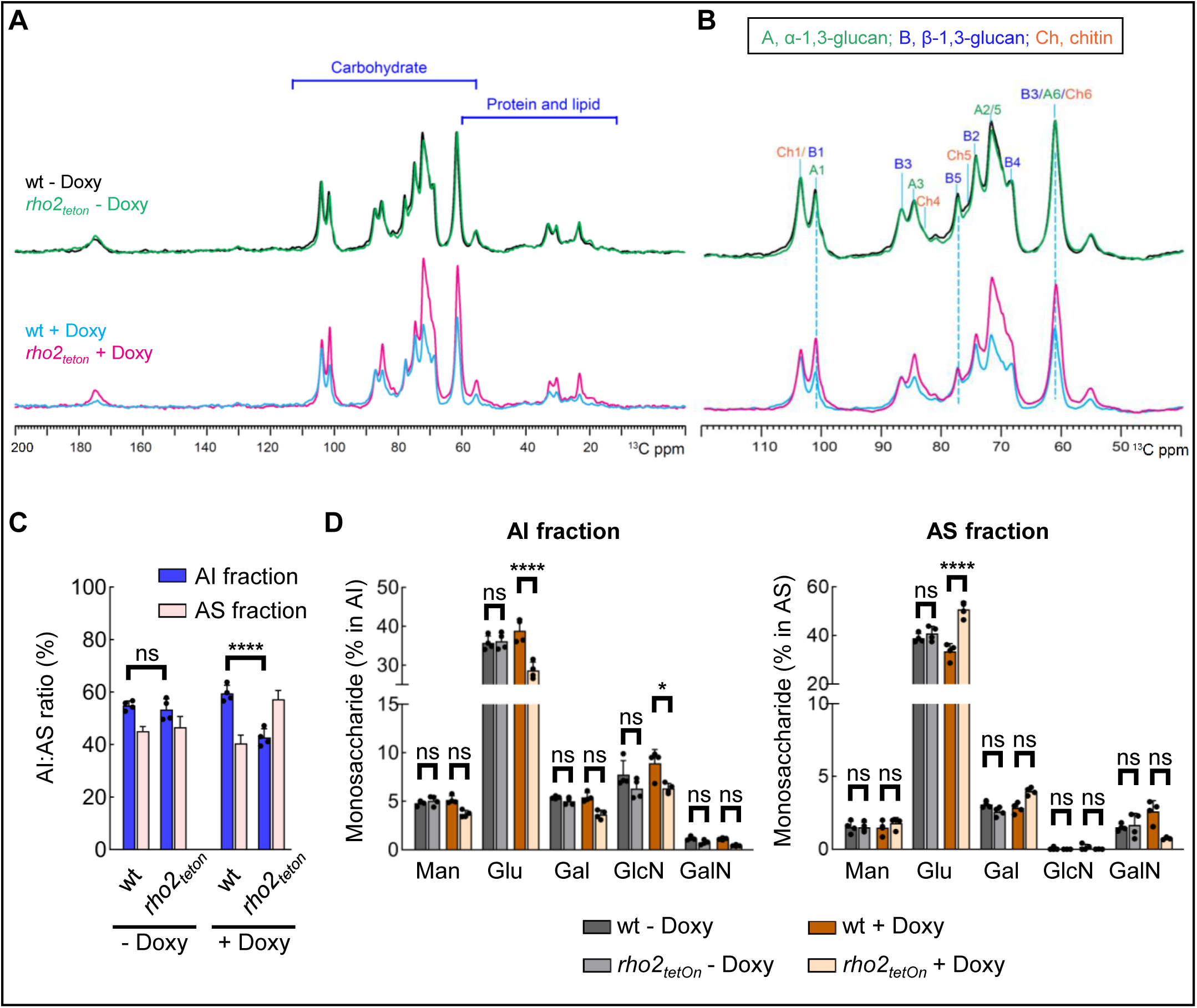
Impact of Rho2 on the cell wall of *A. fumigatus*. (A - D) Conidia of wild type and the conditional *rho2_tetOn_* mutant were inoculated in AMM supplemented with or without 7.5 µg ml^-1^ doxycycline and incubated in a rotary shaker at 37°C. After 23 h incubation, mycelium was harvested and analyzed with ssNMR (A and B) or chemically (C and D). (A and B) Representative 1D ^13^C cross polarization spectra of rigid cell wall molecules obtained for the indicated strains were overlaid for comparison (number of biological replicates: 3; see Supplementary Fig. 4 for all spectra). (A) The representative carbohydrate and protein/lipid regions are marked. (B) Zoom-in view of the carbohydrate regions of the 1D ^13^C spectra shown in (A), with key peaks labeled for different carbon sites of chitin (Ch), α-1,3-glucan (A), and β-1,3-glucan (B). For example, the A1 peak at 101 ppm represents the carbon 1 of α-1,3-glucan. (A and B) The spectra were normalized by β-1,3-glucan B3 (86 ppm) peak. (C) The ratio of alkali-insoluble (AI) and alkali-soluble (AS) fractions in the cell walls of wild-type and *rho2_tetOn_* mutant strains of *A. fumigatus* (number of biological replicates: 3). (D) The monosaccharide composition in the AI and AS fractions were determined by gas-chromatography, upon hydrolysis and derivatization of the AI and AS fractions (number of biological replicates: 3). Man, mannose; Glu, glucose; Gal, galactose, GlcN, glucosamine; GalN, galactosamine. (C and D) Error bars indicate standard deviations. Statistical significance (****, p ≤ 0.0001; *, p ≤ 0.05; ns, not significant) was calculated with the two-way analysis of variance (ANOVA) with Šídák’s multiple comparison post-test.

## DISCUSSION

We and others have previously shown that disruption of central components of the CWI pathway, e.g., Rho1, protein kinase C, or MAP kinases, leads to reduced virulence of fungal pathogens in the genera *Aspergillus*, *Candida* and *Cryptococcus* in murine infection models (Dichtl et al., 2016; Dirr et al., 2010). However, mutations in such central components typically entail pleiotropic effects on the fungal physiology which cause severe growth defects and high susceptibility to multiple forms of stress. It therefore remained unclear whether the reduced virulence is due to a more general growth defect, such as slow growth under harsh conditions, or due to a specific role of the pathway, for example to withstand the immune defense. Defects in central components of the CWI pathway were shown to result in an altered cell wall structure even under normal growth conditions (Dichtl et al., 2016; Gerik et al., 2008; Martín et al., 1993; Navarro-García et al., 1998; Rocha et al., 2015; Román et al., 2016). This could result in a generally weaker cell wall which could increase the fungal sensitivity to the host’s antifungal defense mechanisms or affect the exposure of PAMPs resulting in stronger activation of the host immune system. Furthermore, a dysfunctional CWI pathway could also impair the pathogen’s ability to actively adapt, i.e., optimize and repair the cell wall when it is challenged, e.g., by immune cells.

To unravel this, we first aimed to clarify whether neutrophil granulocytes, which play a key role in the defense against *A. fumigatus*, are able to trigger the cell wall stress response in this fungus. The results obtained with the luciferase-based cell wall stress reporter demonstrate that granulocytes indeed trigger strong activation of the *agsA* promoter, which is a well-characterized target of the CWI pathway in *Aspergillus* species (Rocha et al., 2016). Notably, the activation of the reporter occurred with a delay of 50-60 minutes after adding the granulocytes. This lag is in very good agreement with our previously reported kinetics of the killing activity of granulocytes under similar conditions (Ruf et al., 2018).

Next, we examined the susceptibility of *A. fumigatus* mutants that have an impaired CWI or CWI pathway to killing by granulocytes. Notably, these mutants do not show a severe defect with respect to radial growth under normal growth conditions (Dichtl et al., 2012). The cell wall stress sensor Wsc1, which is important to cope with echinocandin-induced cell wall stress (Dichtl et al., 2012), appears to be dispensable for withstanding killing by granulocytes. In contrast, mutants lacking the stress sensor MidA or the Rho GTPases Rho2 or Rho4 are significantly more susceptible to killing by granulocytes. Rho4 is a major regulator of septum formation (Dichtl et al., 2015; Kwon et al., 2011; Si et al., 2010). We have previously shown that septa improve survival of *Aspergillus* hyphae under cell wall stress conditions by sealing off lysed compartments from the remaining hyphal network (Dichtl et al., 2015; Geißel et al., 2018). The increased susceptibility of the Δ*rho4* mutant can therefore easily be explained with its incapability to form septa.

The exact cellular roles of MidA and Rho2 in *A. fumigatus* have not been explored. The lack of either of them results in increased susceptibility to the cell wall stressors calcofluor white and Congo red, and to increased heat sensitivity (Dichtl et al., 2012). Furthermore, MidA seems to be important for activation of the CWI MAP kinase upon cell wall stress (Dichtl et al., 2012). The complemented Δ*rho2* strain showed significantly better survival than the wild type when exposed to granulocytes. Therefore, we focused our subsequent research on Rho2. As shown with our comprehensive analysis of the complemented Δ*rho2* strain, the improved resistance to killing and calcofluor white is clearly linked to multiple integrations of the circular *rho2* plasmid that was used for complementation. In agreement, the conditional *rho2_tetOn_*mutant under repressed and induced conditions respectively shows a very similar susceptibility and resistance to the cell wall perturbing agent calcofluor white and granulocytes as the Δ*rho2* mutant or the complemented strain (*rho2*).

Results of studies performed on Rho2 homologs in different fungal species suggest that this GTPase has a conserved role in CWI. Deletion of the genes that encode the respective Rho2 homologs in *Aspergillus*, *Schizosaccharomyces pombe*, *Neurospora crassa* and *Colletotrichum gloeosporioides* typically resulted in mutants that were more susceptible to cell wall stress (Dichtl et al., 2012; Kwon et al., 2011; Pérez et al., 2018; Richthammer et al., 2012; Xu et al., 2016). In *S. pombe* and *N. crassa*, it has been shown that the respective Rho2 homologs are upstream activators of protein kinase C which in turn may regulate the CWI MAP kinases (Pérez et al., 2018; Richthammer et al., 2012) and, in *S. pombe*, ɑ-1,3-glucan synthesis (Pérez et al., 2018). However, at least some characteristics of Rho2’s role in CWI diverge in different species. For example, while *Aspergillus* mutants which lack the genes that encode the Rho2 homologs are highly susceptible to calcofluor white (Dichtl et al., 2012; Kwon et al., 2011), a respective *S. pombe* Δ*rho2* mutant gets significantly more resistant to this stressor (Villar-Tajadura et al., 2008).

How does Rho2 contribute to the resistance of *A. fumigatus* to killing by human granulocytes? The conditional mutant that expresses a constitutively active Rho2 (Rho2^G20V^) under induced conditions was helpful to answer this question. Expression of Rho2^G20V^ results in a significant increase of calcofluor white-stainable cell wall chitin. This demonstrates that Rho2 is able to upregulate the cell wall biosynthesis. Our cell wall analysis of the Rho2-overexpressing strain also demonstrates that Rho2 is capable of altering the cell wall. Although the changes observed upon overexpression of the non-constitutively active Rho2 appear more subtle than those observed for *A. fumigatus* mutants lacking the major cell wall polysaccharide synthases (Beauvais et al., 2005; Chakraborty et al., 2021; Dichtl et al., 2015; Muszkieta et al., 2014), an overall remarkable change in favor of the alkali-soluble fraction and ɑ-1,3-glucan linked with relative increase of total chitin and relative decrease of β-1,3-glucan is evident. Interestingly, the conditional *rho2* mutant under repressed conditions showed no altered cell wall. This suggests that under normal growth conditions, Rho2 is not required to build the normal cell wall. Together, this suggests that Rho2 gets actively involved in the defense against attacking granulocytes by upregulating cell wall biosynthesis upon cell wall stress, most likely to repair granulocyte-triggered damage.

Rho2 could stimulate cell wall biosynthesis directly, e.g., by activating cell wall synthases, or transcriptionally by inducing expression of cell wall synthases which are targets of the CWI pathway (Katayama et al., 2015; Rocha et al., 2016). The Rho2 homologs in *S. pombe* and *N. crassa* have been reported to activate protein kinase C (Pérez et al., 2018; Richthammer et al., 2012), which is thought to be the major activator of the MAP kinase module of the CWI pathway (Dichtl et al., 2016). If Rho2 were a direct activator of the CWI pathway, then one would expect a dramatic increase in signaling when Rho2^G20V^ is expressed. Indeed, when inducing expression of the constitutively active Rho2^G20V^ in *A. fumigatus*, an increasing response from the luciferase CWI signaling reporter can be detected with some delay. However, even though the Rho2^G20V^-induced signal steadily increments over time, it is quickly outpaced by the signal caused by azole-induced CWI stress. The finding that expression of *rho2*^G20V^ causes an almost complete growth arrest in less than one hour, while after three hours it still yielded only a luciferase reporter signal less than half of the signal caused by azoles, suggests that the Rho2^G20V^-triggered activation of CWI signaling is rather indirect in nature, e.g., by Rho2 triggering changes in the cell wall which in turn trigger CWI signaling. This is in line with our previous finding where Rho2 was not required to trigger phosphorylation of MpkA (Dichtl et al., 2012) and suggests that Rho2 is not a primary activator of the CWI MAP kinase module in *A. fumigatus*. Moreover, expression of Rho2^G20V^ causes changes in the cell wall in a Δ*mpkA* deletion mutant. This proves that Rho2 can trigger increased cell wall biosynthesis independently of the CWI MAP kinase module.

We found that the expression of Rho2^G20V^ in *A. fumigatus* is lethal. While the Rho2^G20V^-expressing hyphae remain viable for several days, they eventually die. It is intriguing to speculate that this lethal effect could be linked with the evident excessive cell wall biosynthesis, similar as it was previously found for azole-treated *Aspergillus* (Geißel et al., 2018). Our finding that expression of Rho2^G20V^ is not lethal if it occurs in a Δ*mpkA* deletion mutant supports this hypothesis. Due to the presumably lower basal expression of enzymes important for cell wall biosynthesis, the Rho2^G20V^-triggered “toxic” cell wall biosynthesis is less severe, which can be visualized microscopically by the formation of smaller cell wall changes. Interestingly, the excessive Rho2^G20V^-triggered cell wall biosynthesis takes place particularly at certain points of the cell wall, which leads to cell wall bulges. We previously described that *A. fumigatus* treated with azole antifungals also triggers excessive cell wall biosynthesis with the formation of chitin-rich patch-like structures, and we have shown that the excessive cell wall biosynthesis contributes to the fungicidal effect of this class of antifungals (Geißel et al., 2018). Rho2 could therefore be a player in the formation of patches that get formed if the 14α-demethylase (CYP51) is inhibited. However, since the Δ*rho2* mutant also forms cell wall patches after exposure to azoles, Rho2 appears to be not involved in the formation of azole-induced cell wall alterations.

Is Rho2 involved in virulence overall or is it merely an *in vitro* factor that helps hyphae to survive when exposed to granulocytes? A role of Rho2 homologs for the virulence in animal models had not been established for any other fungal pathogens so far. The results of our *Galleria mellonella*-based infection experiment indicate that Rho2 is involved in virulence since the *A. fumigatus* Δ*rho2* deletion mutant is less virulent when compared with the wild type. However, even though the complemented strain is much more resistant to killing by granulocytes, it does not show an increased virulence in the *Galleria mellonella* infection experiment. An explanation for this could be the overall reduced growth rate of *A. fumigatus* upon overexpression of Rho2. While the complemented Δ*rho2* strain is more resistant to killing by immune cells, its growth is reduced, possibly hampering its virulence. Nevertheless, under certain scenarios an advantage of increased expression of Rho2 is conceivable, e.g., in chronic *Aspergillus* infections as this may support persistence of *Aspergillus*.

In summary, although some questions could not be fully elucidated in our study, a clear model for the role of the cell wall integrity pathway as well as Rho2 in virulence and resistance to neutrophil granulocytes could be developed (Fig. 9). Our results show that neutrophil granulocytes cause cell wall stress when they attack hyphae of *A. fumigatus*. Based on its role in cell wall integrity and importance for withstanding neutrophils, MidA could be a key stress sensor for this stress. Rho2, which is localized at the cell membrane, upregulates cell wall biosynthesis, predominantly of ɑ-1,3-glucan and chitin, to actively counteract granulocyte-induced cell wall damage. This Rho2-dependent upregulation is more of direct nature and less through transcriptional upregulation of enzymes important for cell wall biosynthesis. This ability of *Aspergillus* to adapt to the stress caused by neutrophil granulocytes by upregulating cell wall biosynthesis actively supports its virulence.

**Fig. 9.**
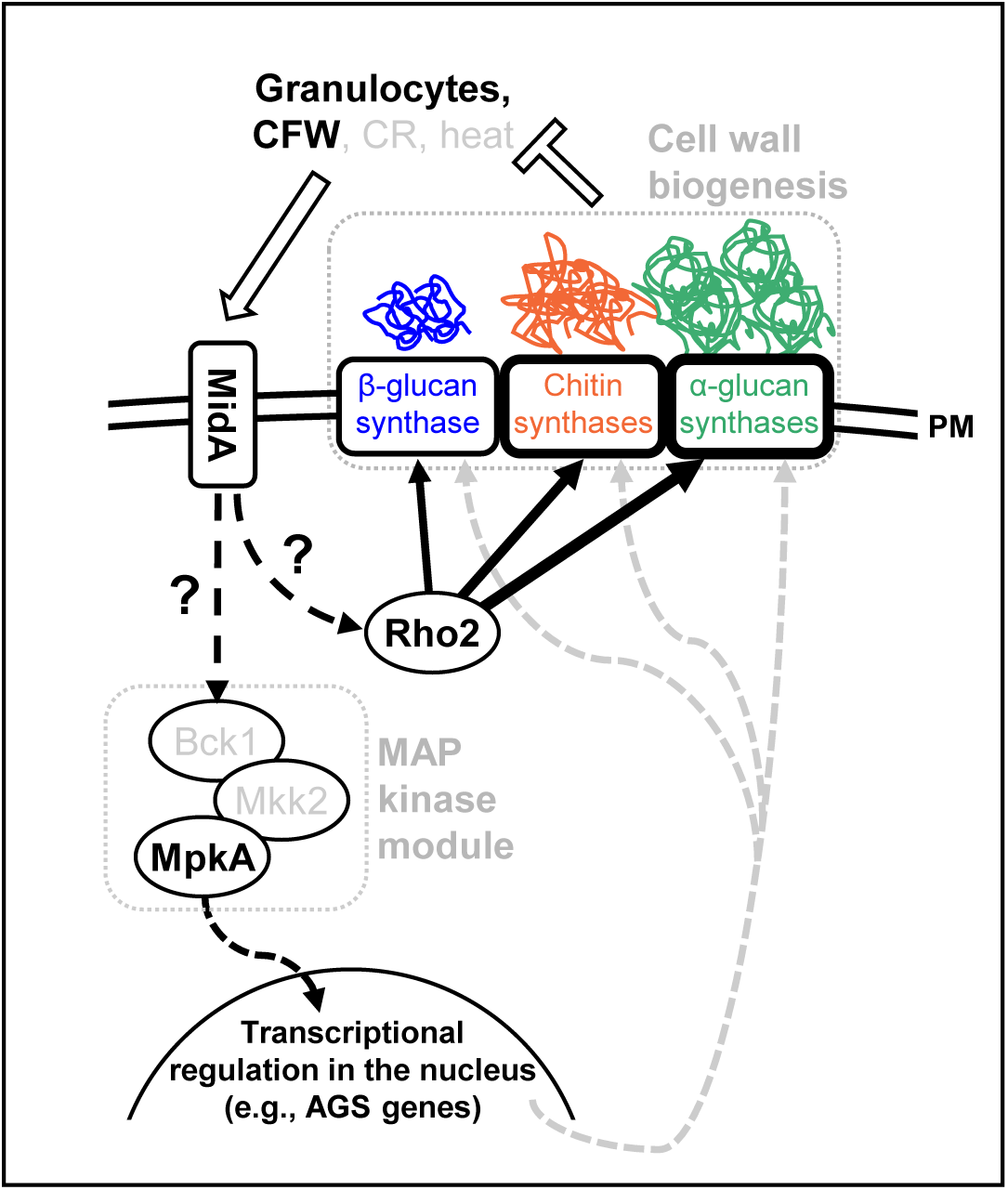
Model of the role of Rho2 in regulation of cell wall biogenesis in *A. fumigatus*. Model of the role of the CWI pathway in withstanding the antifungal activity of granulocytes and supporting virulence. Granulocyte-induced cell wall stress is sensed by the CWI pathway, possibly by the CWI stress sensor MidA. Rho2 triggers a response to this stress, primarily by inducing a general upregulation of cell wall biogenesis. The response involves an upregulation of the biosynthesis of chitin, β-1,3-glucan and other cell wall components (galactomannan, galactosaminogalactan; not shown) and in particular of α-1,3-glucan. The CWI mitogen activated protein (MAP) kinase (K) module consisting of Bck1 (MAPKKK), Mkk2 (MAPKK) and MpkA (MAPK), supports this response, most likely by ensuring appropriate levels of expression of cell wall biosynthetic enzymes. AGS, alpha glucan synthase; PM, plasma membrane.

## MATERIAL AND METHODS

### Strains

The *A. fumigatus* strains used in this study are listed in Table 1. AfS35, a nonhomologous end joining-deficient derivative of D141 (Krappmann et al., 2006; Wagener et al., 2008), served as wild-type strain for all mutants in this work. Mitochondria were visualized with mitochondria-targeted green fluorescent protein (mtGFP) by transforming the respective parental strains with pCH005, essentially as described before (Neubauer et al., 2015). The new plasmid for complementing the Δ*rho2* deletion mutant (pDR003) was constructed by cloning the *rho2* gene including approx. 1.1 kb of its 5’ untranslated region and 1.0 kb of its 3’ untranslated region into pSK379 (a kind gift from Sven Krappmann). pDR003 was subsequently transformed into the Δ*rho2* deletion mutant to obtain the complemented mutants (*rho2_add3’_*) with one (#8-11) or more than one integration (#1-3,5-7) of the plasmid. The wild-type strain expressing the corrected N-terminally GFP-tagged Rho2 was constructed as described before (Dichtl et al., 2012), but using a new 5’ primer to include the full coding sequence of *rho2* to construct the respective expression plasmid (pKD005). The conditional *rho2_tetOn_* mutant was constructed by inserting the doxycycline-inducible *oliC*-tetOn promoter cassette derived from pJW128 before the coding sequence of *rho2* by double-crossover homologous recombination, essentially as described before (Helmschrott et al., 2013). The conditional *rho2*^G20V^*_tetOn_* was constructed in a similar way. However, the 3’ region used for homologous recombination was amplified using a mutated *rho2* allele as template. This mutated *rho2* allele was constructed by site-directed mutagenesis of the plasmid pKD005. The site-directed mutagenesis resulted in deletion of the first intron of *rho2* and in mutation of the codon at position 239 (239GGA>GTA), resulting in an amino acid exchange at position 20 (20G>V) in the encoded protein sequence. The luciferase-based *rho2*^G20V^*_tetOn_* cell wall stress reporter strain *rho2*^G20V^*_tetOn_* + *agsA(p)_A.niger_-Fluc* was constructed by transforming the *rho2*^G20V^*_tetOn_* strain with pBG005-phleo (Geißel et al., 2018). The Δ*mpkA* deletion mutants were constructed using a self-excising hygromycin B resistance cassette (obtained from pSK528), essentially as described before (Hartmann et al., 2010; Wagener et al., 2008).

**Table 1.** *A. fumigatus* strains used in this work.

| Strain or genotype | Relevant genetic modification | Parental strain | Reference |
| --- | --- | --- | --- |
| AfS35 (wt) | *akuA::loxP* | D141 | (Krappmann et al., 2006) |
| wt + sGFP | pJW103 | AfS35 | (Dichtl et al., 2015) |
| wt + *agsA(p)<sub>A.niger</sub>-Fluc* | pBG005-phleo | AfS35 | (Geißel et al., 2018) |
| $\Delta midA$ | *midA::loxP-hygro<sup>R</sup>/tk* | AfS35 | (Dichtl et al., 2012) |
| *midA* | pSK379-*midA(p)-midA* | $\Delta midA$ | (Dichtl et al., 2012) |
| $\Delta wsc1$ | *wsc1::loxP-hygro<sup>R</sup>/tk* | AfS35 | (Dichtl et al., 2012) |
| $\Delta rho2$ | *rho2::loxP-hygro<sup>R</sup>/tk* | AfS35 | (Dichtl et al., 2012) |
| *rho2* | pSK379-*rho2(p)-rho2* | $\Delta rho2$ | (Dichtl et al., 2012) |
| $\Delta rho4$ | *rho4::loxP-hygro<sup>R</sup>/tk* | AfS35 | (Dichtl et al., 2012) |
| *rho4* | pSK379-*rho4(p)-rho4* | $\Delta rho4$ | (Dichtl et al., 2012) |
| AfS35 + mtGFP | pCH005 | AfS35 | (Ruf et al., 2018) |
| $\Delta midA$ + mtGFP | pCH005 | $\Delta midA$ | This work |
| *midA* + mtGFP | pCH005 | *midA* | This work |
| $\Delta wsc1$ + mtGFP | pCH005 | $\Delta wsc1$ | This work |
| $\Delta rho2$ + mtGFP | pCH005 | $\Delta rho2$ | This work |
| *rho2* + mtGFP | pCH005 | *rho2* | This work |
| $\Delta rho4$ + mtGFP | pCH005 | $\Delta rho4$ | This work |
| *rho4* + mtGFP | pCH005 | *rho4* | This work |
| *rho2<sub>add3'</sub>* | pDR003 | $\Delta rho2$ | This work |
| *rho2<sub>tetOn</sub>* | *rho2(p)::ptrA-tetOn* | AfS35 | This work |
| wt + GFP-Rho2 | pKD005 | AfS35 | This work |
| *rho2<sub>tetOn</sub> ΔmpkA* | *mpkA::loxP-hygro<sup>R</sup>/blaster* | *rho2<sub>tetOn</sub>* | This work |
| *rho2<sub>tetOn</sub>* + mtGFP | pCH005 | *rho2<sub>tetOn</sub>* | This work |
| *rho2<sup>G20V</sup><sub>tetOn</sub>* | *rho2(p)::ptrA-tetOn; rho2<sup>G20V</sup>* | AfS35 | This work |
| *rho2<sup>G20V</sup><sub>tetOn</sub>* + mtGFP | pCH005 | *rho2<sup>G20V</sup><sub>tetOn</sub>* | This work |
| *rho2<sup>G20V</sup><sub>tetOn</sub>* + *agsA(p)<sub>A.niger</sub>-Fluc* | pBG005-phleo | *rho2<sup>G20V</sup><sub>tetOn</sub>* | This work |
| *rho2<sup>G20V</sup><sub>tetOn</sub> ΔmpkA* | *mpkA::loxP-hygro<sup>R</sup>/blaster* | *rho2<sup>G20V</sup><sub>tetOn</sub>* | This work |
| $\Delta mpkA$ | *mpkA::loxP-hygro<sup>R</sup>/blaster* | AfS35 | This work |

### Culture Conditions and Chemicals

Experiments were performed either in or on *Aspergillus* minimal medium (AMM) (Hill and Kafer, 2001), RPMI 1640 medium (with L-glutamine, but without phenol red; 11835-063; Gibco, Thermo Fisher, Waltham, MA, USA), RPMI 1640/PR (with L-glutamine and with phenol red; R8758; Sigma-Aldrich, St. Louis, MO, USA), RPMI 1640/MOPS medium (RPMI 1640 medium (with L-glutamine, but without bicarbonate and without phenol red; R8755; Sigma-Aldrich) supplemented with glucose to a final concentration of 2 % (w/v) and with 3-(N-morpholino) propanesulfonic acid (MOPS) at a final concentration of 0.165 mol/L, pH 7.0), yeast extract glucose medium (YG; 2 % (w/v) D-glucose; 0,5 % (w/v) yeast extract (); pH 6.0), or Sabouraud medium (Sab; 4% (w/v) D-glucose, 1% (w/v) peptone (#LP0034; Thermo Fisher Scientific; Rockford, IL, USA); pH 7.0 ± 0.2). Solid media were supplemented with 2% (w/v) agar (214030; BD Bioscience, Heidelberg, Germany). All strains were maintained on AMM to harvest conidia. Experiments performed in RPMI 1640 were incubated with 5% CO_2_. Paraformaldehyde (158127), calcofluor white (F3543) and Congo red (60910) were obtained from Sigma-Aldrich. Doxycycline was obtained from Clontech (631311; Mountain View, CA, USA). Percoll was obtained from GE Healthcare (10253000; Uppsala, Sweden). Luciferin was purchased from Promega (E1601; Fitchburg, WI, USA). Lipopolysaccharides (LPS) were obtained from Invivogen (tlrl-peklps; San Diego, CA, USA).

### Cell wall stress reporter assay

To analyze activation of the CWI pathway by human granulocytes, approximately 2 x 10^5^ conidia of the luciferase-based *A. fumigatus* cell wall stress reporter strain wt + *agsA(p)_A.niger_-Fluc* (Geißel et al., 2018) were inoculated in RPMI 1640 medium per well in white 96-well polystyrene microplates with transparent bottom (#655095, #656171) purchased from Greiner Bio-One (Kremsmünster, Austria). After 10 h incubation at 37°C with 5% CO_2_, medium was supplemented with 0.5 mM luciferin and, when indicated, with the indicated amount of calcofluor white, or the indicated number of human granulocytes or A549 cells (86012804-1VL, Merck, Darmstadt, Germany) was added. The microplates were then incubated at 37°C with 5% CO_2_ and the luminescence was measured over time from the bottom with a Clariostar microplate reader (BMG Labtech; Ortenberg, Germany).

To analyze activation of the CWI pathway by Rho2^G20V^, approximately 2 x 10^4^ conidia of the luciferase-based cell wall stress reporter strain *rho2*^G20V^*_tetOn_* + *agsA(p)_A.niger_-Fluc* were inoculated in RPMI 1640/MOPS medium per well in white 96-well polystyrene microplates without transparent bottom (#3917) purchased from Corning Inc. (Corning, NY, USA). After 7 h incubation at 37°C, medium was supplemented with 0.5 mM luciferin and, when indicated, with the indicated amount of calcofluor white or doxycycline was added. The microplates were then incubated at 37°C and the luminescence was measured over time from the top with a GloMax Explorer microplate reader (Promega).

### MitoFLARE assay with human granulocytes

The killing activity of human granulocytes against *A. fumigatus* hyphae was analyzed following our previously described protocol (Brantl et al., 2021; Ruf et al., 2018). 3 × 10^3^ conidia of strains that express mtGFP were inoculated in 300 µl RPMI 1640 per well in µ-Slide 8-well microscope slides (80826; Ibidi, Martinsried, Germany) and incubated at 37°C with 5% CO_2_. When indicated, medium was additionally supplemented with doxycycline or 10% (v/v) human serum. After 10 h incubation, 1.5 × 10^6^ granulocytes resuspended in 100 µl RPMI 1640 were added to each well. After 2 h incubation with granulocytes, samples were fixed with 4% (w/v) paraformaldehyde for 10 min. Samples were then stained with calcofluor white (1mg ml^-1^ in double-distilled water) for 10 min followed by washing with phosphate-buffered saline (PBS). Samples were subsequently analyzed with a fluorescence microscope and a 63x objective with oil immersion as described before (Ruf et al., 2018). Different strains and conditions were randomly located in each m-Slide 8-well slide and the assessor was blind to sample identity. In each experiment, 60 hyphae per sample and three samples per condition were analyzed. Prior to the analysis, the killing efficacy of the batch of isolated granulocytes was evaluated with a nonblind wild-type control sample. With the exception of the experiment with serum, experiments in which excessive or no significant killing was found (hyphal vitality of wild type after killing for 2 h of <30% or >90%) were excluded and not considered in the subsequent statistical analysis: Of 21 experiments, one was excluded because of too-high killing activity and two experiments because of too-low killing activity. Statistical significance was calculated with the one-way analysis of variance (ANOVA) with Tukey’s multiple comparison post-test when comparing more than two groups or a two-tailed unpaired (assuming unequal variances) Student’s *t*-test when comparing two groups using GraphPad Prism 5 (GraphPad Software, La Jolla, CA, USA).

### Isolation of human granulocytes

Granulocytes were isolated from the blood of healthy adult volunteers as described before (Ruf et al., 2018). Collection of blood was conducted according to the Declaration of Helsinki and the study was approved by the Ethics Committee of LMU Munich (Reg. Nr. 519-95). Volunteers gave informed written consent.

### MitoFLARE assay after expression of *rho2*^G20V^

Conidia of the *rho2*^G20V^*_tetOn_* expressing mtGFP were inoculated in 200 µl AMM per well in µ-Slide 8-well microscope slides and incubated at 37°C. After 8 h, 100 µl AMM supplemented with doxycycline was added, resulting in a final concentration of 7.5 µg ml^-1^ doxycycline in the well. The slides were then incubated at 37°C and analyzed at the indicated time points with a spinning disc confocal fluorescence microscope similar as described before (Brantl et al., 2021). At least 50 hyphae were analyzed at each time point.

### Microscopy

Growth and germination of mutants was assessed microscopically with a Lionheart FX automated microscope (Agilent BioTek; Santa Clara, CA, USA). Quantitative analysis of the killing activity of granulocytes against *A. fumigatus* hyphae with MitoFLARE strains was performed with a Leica DM IRB inverted fluorescence microscope (Leica Microsystems, Mannheim, Germany). Imaging and quantitative analysis of the viability of the *rho2*^G20V^*_tetOn_* MitoFLARE strain and imaging of calcofluor white-stained hyphae (3.33 µg ml^-1^ calcofluor white, followed by after washing) after *rho2*^G20V^ expression or voriconazole treatment was performed with an Eclipse Ti2 confocal microscope (Nikon; Tokio, Japan) equipped with a CSU-W1 spinning disc confocal scanner unit (Yokogawa; Tokio, Japan) equipped with a climate chamber. Imaging of the subcellular localisation of GFP-tagged Rho2 was performed with a Leica SP5 inverted confocal laser scanning microscope (Leica Microsystems, Mannheim, Germany) equipped with a climate chamber.

### Bioinformatics

Sequences were obtained from FungiDB (Amos et al., 2022) and alignments were performed with Jalview (Waterhouse et al., 2009). Plasmids were planned and analyzed with Benchling (https://benchling.com).

### Galleria mellonella infection experiments

Infection experiments were performed as described previously (Geißel et al., 2017; Neubauer et al., 2015). For each condition, the survival of 60 larvae (obtained from AKM − Angel- und Ködermarkt, Munich, Germany) was analyzed. Experiments were performed in groups of 20 larvae for each condition. The larvae were infected with 5 × 10^5^ conidia in 10 µl ddH_2_O per larva. Control groups were either not injected or injected with 10 µl ddH_2_O. The larvae were then incubated in the dark at 37°C and viability was assessed every 8 h. Survival was estimated with a Kaplan-Meier curve. Significance was calculated with GraphPad Prism 5 using a log-rank (Mantel-Cox) test.

### Cell wall analysis

Conidia of the strains were inoculated in AMM supplemented with or without 7.5 µg ml^-1^ doxycycline and incubated in a rotary shaker (180 rpm) at 37°C. For the solid-state NMR (ssNMR) analysis, mycelium was harvested after 23 h, washed with AMM, frozen, and subjected to analysis. The 1D high-resolution ssNMR analyses were conducted on a Bruker Advance 600 MHz (14 Tesla) spectrometer using 3.2 mm magic-angle spinning (MAS) HCN probes under 15 kHz MAS at 298 K. The ^13^C chemical shifts were externally referenced to the ^13^C signal of the adamantane CH_2_ group at 38.48 ppm on the tetramethylsilane (TMS) scale. The magic angle was calibrated using KBr. Typical radiofrequency field strengths were 62.5-100 kHz for ^1^H and 50-62.5 kHz for ^13^C. The initial magnetization for the experiment was created using ^1^H-^13^C cross-polarization that preferentially detects rigid molecules. Typically, 1 ms Hartmann-Hahn contact was used for the cross polarization (CP).

For chemical analysis of the cell wall, mycelium was harvested after 23 h, washed with MilliQ water, and subjected to analysis. After fractionating the cell wall polysaccharides into alkali-soluble and insoluble fractions, they were acid-hydrolyzed, derivatized and subjected to gas-chromatography analysis to determine their monosaccharide composition, following the protocol described earlier (Richie et al., 2009). The analysis was performed on three biological replicates of the cell wall alkali-soluble and insoluble fractions extracted. Statistical significance was calculated with the two-way analysis of variance (ANOVA) with Šídák’s multiple comparison post-test using GraphPad Prism 10 (GraphPad Software).

## AUTHOR CONTRIBUTIONS

J.W. conceived the study. J.W., D.R. and K.S. wrote the manuscript draft. T.W. and V.A. reviewed and edited the manuscript. K.S., L.D.F. and T.W. performed the solid-state NMR cell wall analysis of *Aspergillus* mycelium. M.L and V.K. performed the chemical cell wall analysis. All other experiments and analyses were planned and performed by D.R., K.S., S.B., H.E., V.B., P.H., K.D., and J.W. All authors read and authorized the manuscript.

## ACKNOWLEDGMENTS

This work was in part supported by the German Research Foundation (430055013; J.W.), the Department of Clinical Microbiology, School of Medicine, Trinity College Dublin, the Graduate School of Life Sciences (GSLS) at the University of Würzburg and the Förderprogramm für Forschung und Lehre (FöFoLe) of the Medical Faculty of the Ludwig-Maximilians-Universität München. The ssNMR characterization by K.S., L.D.F. and T.W. was supported by the National Institutes of Health (NIH) AI173270. The chemical analysis of the cell wall by M.L and V.A. was supported by ANR-10-LABX-62-IBEID and ANR-21-CE17-0032-01; FUNPOLYVAC grants.

## DECLARATION OF THE USE OF GENERATIVE ARTIFICIAL INTELLIGENCE

During the preparation of this work the authors used DeepL (deepl.com; Cologne, Germany) for language optimization, i.e., to identify synonyms and to correct errors in spelling, grammar, and punctuation.

## FIGURE LEGENDS

**Supplementary fig. 1.**
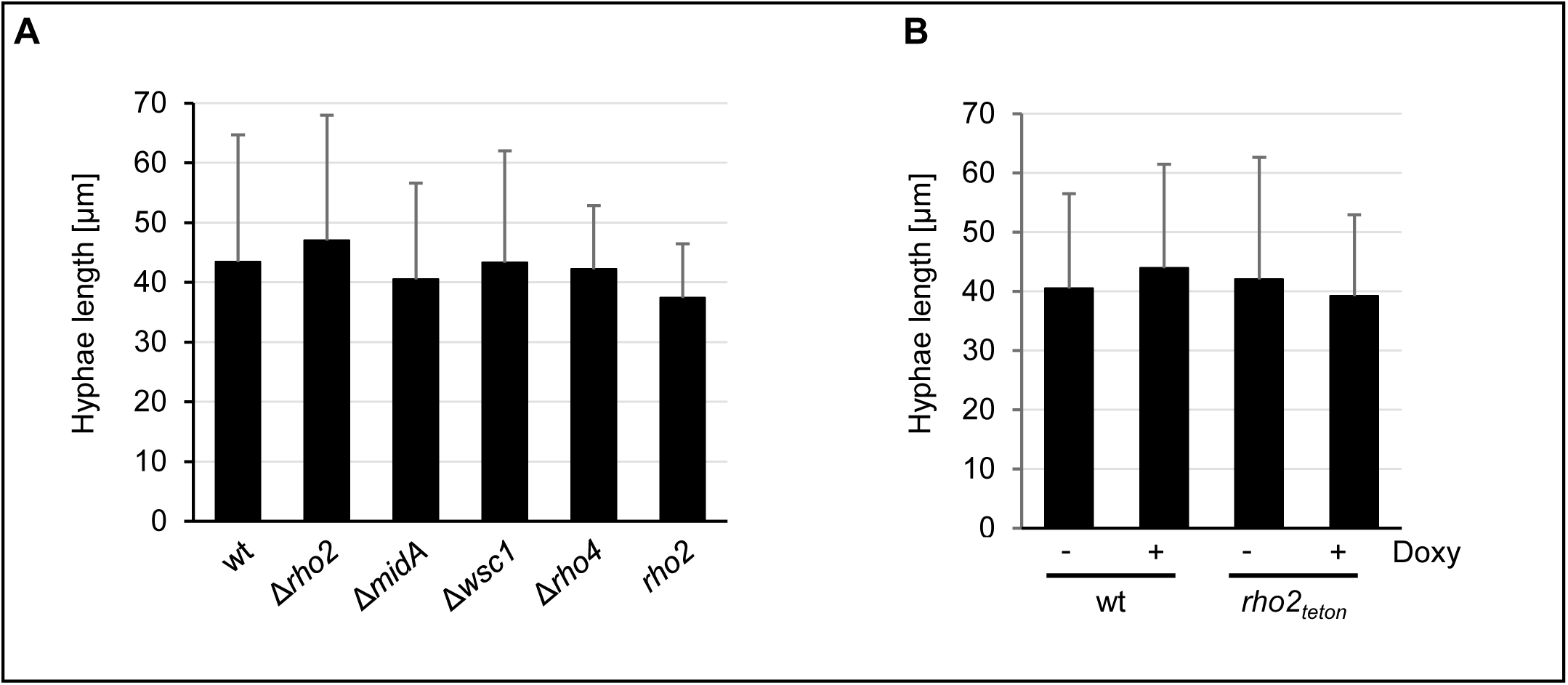
Growth rates of germinating conidia of *A. fumigatus* cell wall integrity mutants used in this study. (A and B) Conidia of the indicated strains were inoculated in RPMI-1640 (with phenol red; RPMI-1640/PR) and incubated at 37°C with 5% CO_2_. After 9.5 h incubation, the length of hyphae (n=28 per strain and condition) was microscopically assessed and the mean lengths were plotted in the graph. Germination rate was >98 % for all strains. The error bars indicate standard deviations.

**Supplementary fig. 2.**
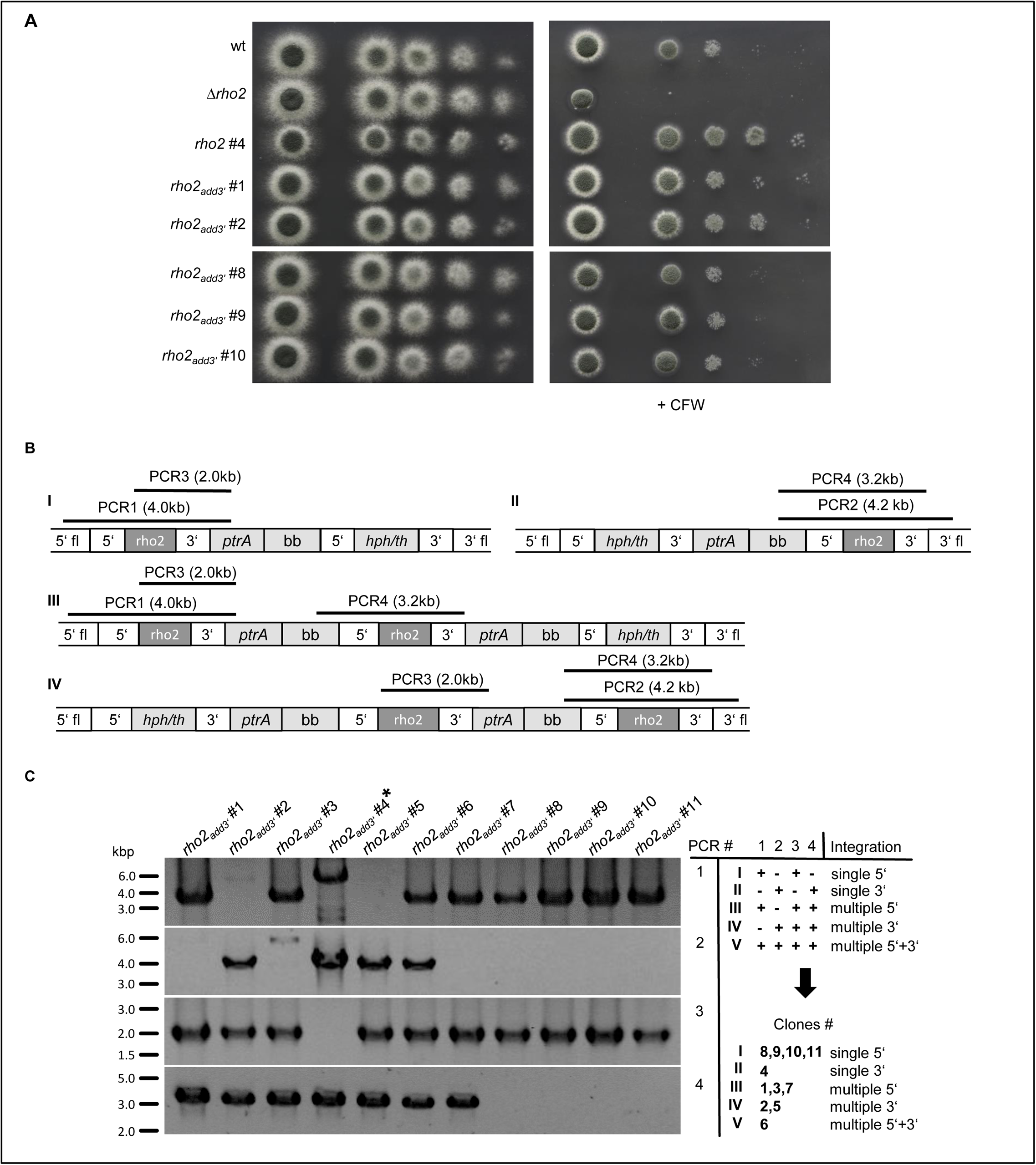
Analysis of *A. fumigatus* mutants with single or multiple integrations of the *rho2* gene. (A) In a series of 10-fold dilutions derived from a starting suspension of 5 × 10^7^ conidia ml^-1^ of the indicated strains, aliquots of 3 µl were spotted on AMM agar plates. The numbers (#) indicates the respective clone number. When indicated, medium was supplemented with calcofluor white (CFW). Representative images were taken after 34 h incubation at 37°C. (B and C) Diagnostic PCRs were performed to determine the number of integrations of the *rho2* plasmid (pDR003) used to complement the Δ*rho2* deletion mutant. (B) The coding region of *rho2* was deleted with a hygromycin resistance cassette (hph/tk). The complementation plasmid harbors the *rho2* gene with its 5’ and 3’ flanking (fl) region as well as the plasmid backbone (bb) and the pyrithiamine resistance gene. This allowed for integration of the plasmid by homologous recombination either at the 5’ or 3’ flanking region at the *rho2* locus in the chromosome. A single integration of the plasmid at the 5’ or 3’ flanking region is shown in I and II, respectively. Sequential integrations of the plasmid at the 5’ or 3’ flanking region are shown in III and IV, respectively. Overlapping PCRs were performed to reveal the integration of the plasmid at the 5’ flanking region (PCR1) or 3’ flanking region (PCR2). PCR3 and PCR4 were performed to reveal sequential integrations of the plasmids. The expected fragment sizes for the PCRs are shown in brackets. (C) Multiple clones of the Δ*rho2* mutant that was transformed with pDR003 (*rho2_add3’_* #1-11) were analyzed. The results for the indicated PCRs are shown on the left. *rho2_add3’_* #4 (labeled with an asterisk) yielded a PCR product of the wrong size in PCR1 and was therefore excluded from subsequent analyses. An interpretation of the number of integrations of the plasmid at the respective sites are shown on the right. kbp, kilobase pairs.

**Supplementary fig. 3.**
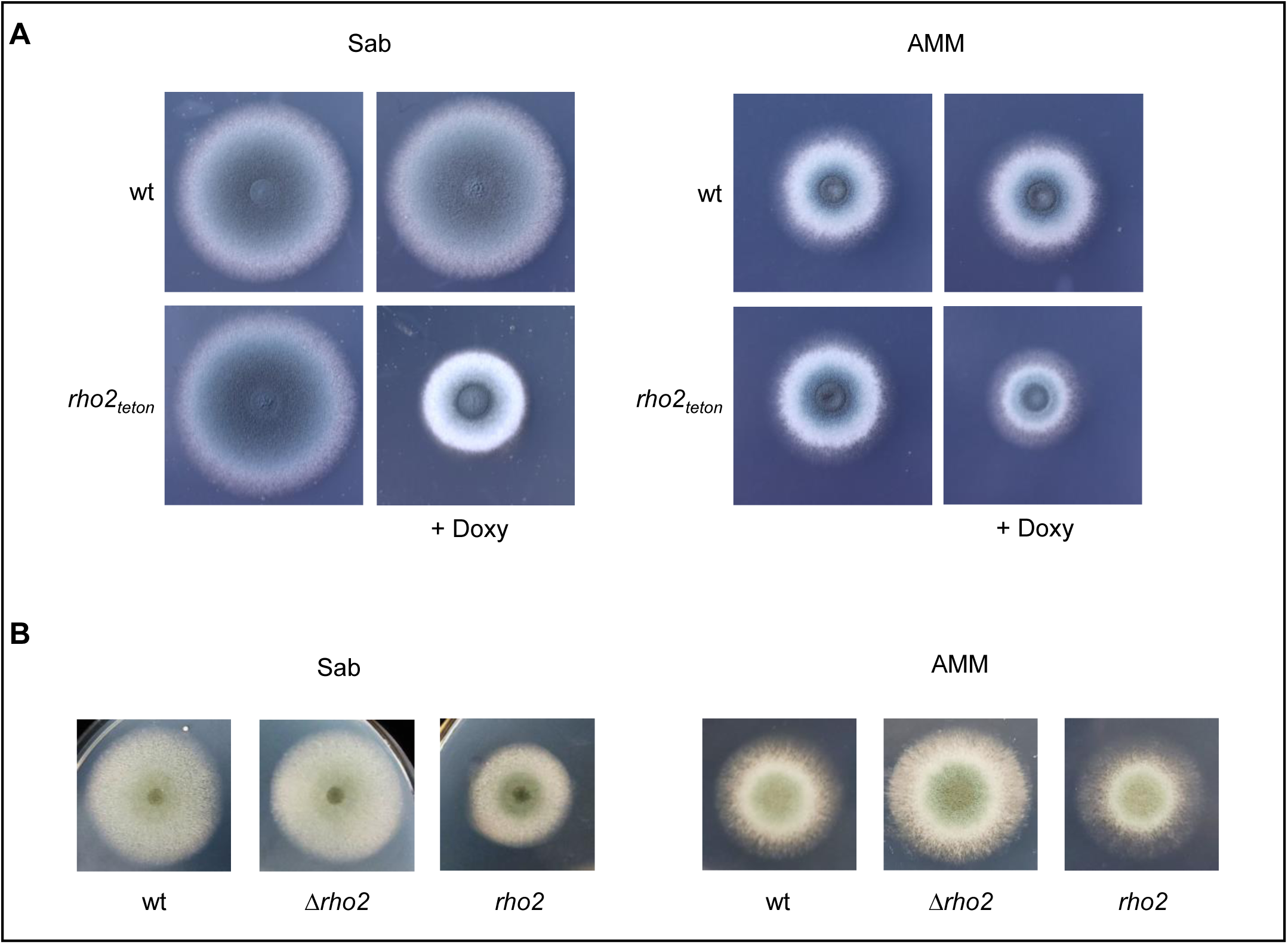
Impact of *rho2* overexpression on radial growth. (A and B) 1.5 × 10^5^ conidia of the indicated strains were spotted on Sabouraud (Sab) or AMM agar plates in triplicates. When indicated, medium was supplemented with 7.5 µg ml^-1^ doxycycline (+Doxy). Representative images were taken after 46 h incubation at 37°C.

**Supplementary fig. 4.**
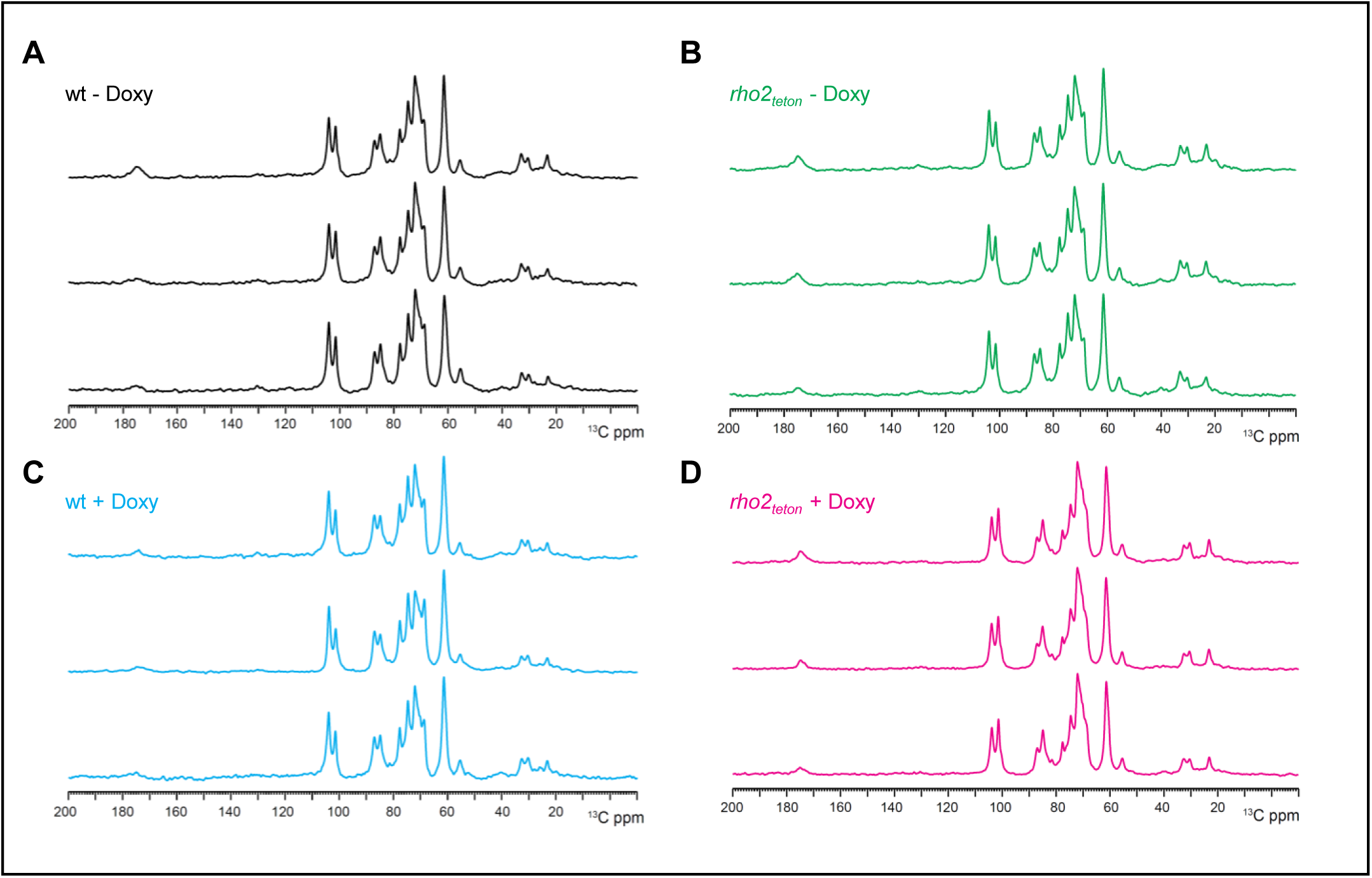
Solid-state NMR 1D ^13^C cross polarization spectra of three biological replicates of the indicated strains cultured with or without doxycycline. Conidia of wild type and the conditional *rho2_tetOn_* mutant were inoculated in AMM supplemented with or without 7.5 µg ml^-1^ doxycycline and incubated in a rotary shaker at 37°C. After 23 h incubation, mycelium was harvested and analyzed with ssNMR. 1D ^13^C cross polarization spectra obtained for three biological replicates of the indicated strains are depicted. See Figure 8 for overlays of representative spectra.

**Supplementary video 1. Expression of Rho^G20V^ results in growth arrest (induced condition).** Conidia of the conditional *rho2*^G20V^*_tetOn_* mutant were inoculated in a 96-well plate in RPMI 1640/MOPS medium. After 7 h incubation at 37°C, medium was supplemented with 25 µg ml^-1^ doxycycline and growth was monitored with an automated microscope over time.

**Supplementary video 2. Expression of Rho^G20V^ results in growth arrest (non-induced condition; control).** Conidia of the conditional *rho2*^G20V^*_tetOn_* mutant were inoculated in a 96-well plate in RPMI 1640/MOPS medium (control). After 7 h incubation at 37°C, growth was monitored with an automated microscope over time.

